# Distinct Regulation of Bioenergetics and Translation by Group I mGluR and NMDAR

**DOI:** 10.1101/552638

**Authors:** Sudhriti Ghosh Dastidar, Shreya Das Sharma, Sumita Chakraborty, Sumantra Chattarji, Aditi Bhattacharya, Ravi S Muddashetty

## Abstract

Neuronal activity is responsible for large energy consumption within the brain. However, the cellular mechanisms draining ATP upon the arrival of a stimulus are yet to be explored systematically at the post-synapse. Here we provide evidence that a significant fraction of ATP is consumed upon glutamate stimulation to energize the mGluR-induced protein synthesis. We find that both mGluR and NMDAR alter protein synthesis and ATP consumption with distinct kinetics at the synaptic-dendritic compartments. While mGluR activation leads to a rapid and sustained reduction in the neuronal ATP level, NMDAR activation has no immediate impact on the same. ATP consumption correlates inversely to the kinetics of protein synthesis for both the receptors. We observe a persistent elevation in protein synthesis within 5 minutes of mGluR activation and robust inhibition of the same within 2 minutes of NMDAR activation, assessed by the phosphorylation status of eEF2 and metabolic labeling. However, a delayed protein synthesis-dependent ATP expenditure ensues after 15 minutes of NMDAR activation. We identify a central role for AMPK in this correlation between protein synthesis and ATP consumption. AMPK is dephosphorylated and inhibited upon mGluR activation while it was rapidly phosphorylated upon NMDAR activation. Perturbing AMPK activity disrupts the receptor-specific modulations of eEF2 phosphorylation and protein synthesis. Therefore, our observations suggest that the glutamate receptors required modulating the AMPK-eEF2 signaling axis to alter neuronal protein synthesis and bioenergetics.

**Short Summary:** Stimulation of glutamate receptors induces robust protein synthesis within cortical neurons and consumes a significantly large fraction of cellular ATP. Glutamate receptors viz. mGlulR and NMDAR modulate AMPK-eEF2 signaling uniquely leading to the dynamic regulation of protein synthesis and bioenergetics.

**Key Highlights:**

- Protein synthesis following glutamate receptor activation is responsible for the bulk of the activity-induced ATP consumption in cortical neurons.
- mGluR and NMDAR regulate protein synthesis with distinct kinetics and dictate the subsequent impacts over neuronal ATP level.
- Dynamic modulation of AMPK and eEF2 phosphorylation is key to create unique temporal features of receptor-specific protein synthesis and bioenergetics.

## Introduction

The brain produces a significant energy burden within the body, even at the ‘resting’ state [1]. Further, the consumption of glucose or oxygen increases with brain activation [2,3]. Within the brain, synapses are the sites of this ATP consumption primarily [4,5] as several synaptic mechanisms are thought to give rise to an exaggerated energy demand within a neuron [5]. Categorizing such mechanisms in terms of their metabolic cost, however, has been difficult due to a lack of conclusive evidence. Earlier studies based on theoretical calculations predicted that the largest amount of ATP is expended to reestablish the ionic gradients following a neuronal spike [5,6]. More recent work, however, demonstrated that the neuronal bursting is quite energy-efficient [7]. Besides, the impact of the synaptic activity on the local energy levels can be quite diverse as the energy supply may vary largely between various neuronal compartments [8]. For example, recent reports have suggested that the rapid endocytosis following glutamate release imposes the highest energy burden at the pre-synaptic nerve terminals [9,10]. Besides, neuronal stimulation induces abundant protein synthesis [11–13] the energy budget of which is still uncharted [14]. Protein synthesis is a determining resource for long-term synaptic plasticity [15–17]. It is an energetically expensive procedure with a requirement of 4 ATP molecules for each round of amino acid incorporation [14,18]. Activation of glutamate receptors such as group I metabotropic glutamate receptors (mGluR) and NMDA Receptors (NMDAR) are reported to alter the rate of protein synthesis [19–21] and are widely implicated to facilitate or induce various forms of synaptic plasticity across different brain regions [22–24].

Neurons meet the enhanced energy demand of activity by inducing ATP production concomitantly. Activity-induced glycolysis, oxidative phosphorylation and the use of glial metabolites like lactate for energy production is key to neuronal function and survival [9, 13, 25–27]. Yet, the link between the activity-induced protein synthesis and energy homeostasis has remained unclear. A recent study, however, pointed out that regulating AMP-activated protein kinase (AMPK) function is critical in maintaining the synaptic ATP balance [25]. Considering the ability of AMPK to act as a metabolic sensor, its competence to alter the catabolic-anabolic balance of the cell [28] and its influence over various forms of synaptic plasticity [29,30], we predicted AMPK to represent the missing link between protein synthesis and energy homeostasis. To test the hypothesis, we asked the following questions: 1) How is the energy level altered on neuronal stimulation? 2) How much of the consumed energy is allocated for protein-synthesis? 3) How do the individual glutamate receptor subtypes generate specific translation responses? And 4) How is the AMP Kinase activity regulated to coordinate the translation and energy supply? Our observations suggest glutamate stimulation in cortical neurons induces robust energy consumption due to the activation of abundant protein synthesis. Both mGluR and NMDAR hold the ability to modulate the AMPK-eEF2K-eEF2 signaling pathway to alter the kinetics of protein synthesis.

## Results

### Protein Synthesis Results in a Significant Metabolic Burden Following Glutamate Stimulation in Cortical Neurons

To assess the impact of protein synthesis on cellular energy content, we stimulated high-density rat cultured cortical neurons with glutamate (25μM) for 5 minutes in the presence or absence of protein synthesis inhibitors (anisomycin or cycloheximide) and quantified the ATP/(ATP+ADP) ratio from cell lysates (**Figure 1A**). Glutamate stimulation led to a sharp drop in the neuronal ATP/(ATP+ADP) ratio, a bulk of which was recovered significantly by pre-incubation with anisomycin (25μM) or cycloheximide (350μM) (**Figure 1B** and **Figure EV 1D**) suggesting a robust ATP consumption due to protein synthesis following the activation of glutamate receptors. To identify the receptors responsible for protein synthesis regulation, we repeated the glutamate stimulation in the presence or absence of D-AP5 (25μM) + CNQX (40μM) (a combination of NMDA and AMPA receptor antagonists) or MPEP (10μM) (mGluR5 antagonist). D-AP5 + CNQX treatment did not rescue the ATP levels unless combined with anisomycin (**Figure 1C**). MPEP pre-treatment, however, significantly rescued the energy drop following glutamate application (**Figure 1C**). To verify this observation further, we stimulated cortical neurons with mGluR and NMDAR specific agonists s-3,5 DHPG (50µM) and NMDA (20µM) respectively **(Figure EV 1A)** and measured the ATP consumption. While DHPG addition led to a significant reduction in the ATP/ATP+ADP ratio, NMDA treatment led to a modest yet not significant reduction in the ATP level (p=0.1501), indicating that the mGluR activity is primarily responsible for protein synthesis-dependent energy consumption on glutamate stimulation.

**Figure 1.**
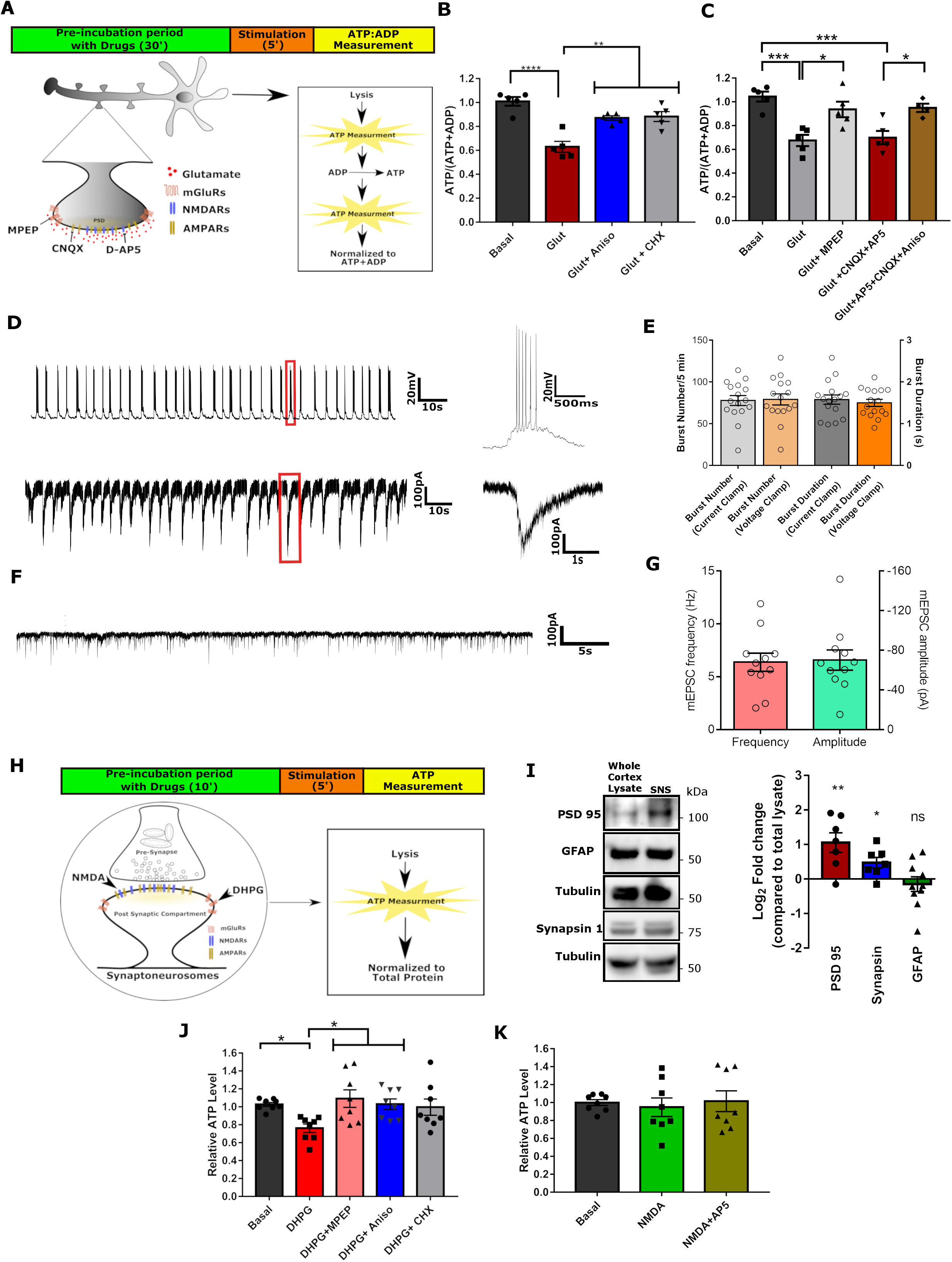
mGluR Dependent Protein Synthesis Presents the Bulk of the Metabolic Burden Following Glutamate Stimulation in Cortical Neurons: **(A)** Schematic depicting experimental workflow for the measurement of the ATP/ATP+ADP ratio using luciferase-based methods following glutamate stimulation in the presence or absence of various drugs from high-density cultured cortical neurons at 15 days *in-vitro* (DIV 15). **(B)** Bar graph representing the normalized average value of the neuronal ATP/ATP+ADP ratio on the basal condition, on glutamate stimulation for 5 minutes, on glutamate stimulation with anisomycin (25μM) and on glutamate stimulation with cycloheximide (350μM). Data presented as mean ± SEM with scattered data points. Values in all other groups were normalized to the basal group for the corresponding experiment. **p<0.01, ***p<0.001, ****p<0.0001, n=5 independent platings. One-way ANOVA followed by Bonferroni’s multiple comparison test. **(C)** Bar graph representing the normalized average value of neuronal ATP/ATP+ADP ratio on the basal condition, on glutamate stimulation for 5 minutes, on glutamate stimulation with MPEP, on glutamate stimulation with CNQX + D-AP5 and on glutamate stimulation with CNQX + D-AP5 + anisomycin. Data presented as mean ± SEM with scattered data points. Values in all other groups were normalized to the basal group for the corresponding experiment.. *p<0.05, ***p<0.001, n=4-5 independent platings. One-way ANOVA followed by Bonferroni’s multiple comparison test. **(D)** Representative current-clamp and voltage-clamp traces showing a single neuron firing spontaneous bursts of action potentials. The traces on the right shows a single representative burst at a higher temporal resolution. **(E)** Bar graph representing the burst frequency and burst duration both in the voltage and current-clamp mode. n=16 neurons from ≥3 independent platings. Data presented as mean ± SEM with scattered data points. **(F)** Representative voltage-clamp traces of miniature EPSCs recorded (V_hold_= - 70mV) from a neuron in presence of TTX. **(G)** Bar graph representing the baseline mEPSC frequency and amplitude in the current-clamp mode. N=11 neurons from ≥3 independent platings. Data presented as mean ± SEM with scattered data points. **(H)** Schematic depicting experimental workflow for the measurement of the synaptic ATP level using luciferase-based methods following DHPG and NMDA treatment in the presence or absence of various drugs from cortical synaptoneurosomes obtained from Sprague Dawley rats at postnatal day 30 (P30). **(I)** Immunoblots depicting levels of PSD 95, GFAP, Synapsin 1 and tubulin in the whole cortical lysate and the synaptoneurosome preparations. Quantification representing the average fold enrichments of PSD 95, Synapsin 1 and GFAP in the synaptoneurosome preparations compared to the whole cortical lysate. Data presented as mean ± SEM with scattered data points. Data points for all the proteins were normalized to their respective whole lysate values. **p<0.05, **p<0.01, n ≥ 7 animals for each group. One-sample t-test. **(J)** Bar graph representing the normalized average value of synaptic ATP level on the basal condition, on DHPG treatment for 5 minutes, on DHPG treatment with MPEP, on DHPG treatment with anisomycin and on DHPG treatment with cycloheximide (350μM). Data presented as mean ± SEM with scattered data points. Values in all other groups were normalized to the basal group for the corresponding experiment. *p<0.05, n=8 animals. One-way ANOVA followed by Bonferroni’s multiple comparison test. **(K)** Bar graph representing the normalized average value of the synaptic ATP level on the basal condition, on NMDA treatment for 5 minutes and on NMDA treatment with D-AP5. Data presented as mean +/- SEM with scattered data points. Values in all other groups were normalized to the basal group for the corresponding experiment. n=8 independent platings.

Since an optimal response to a stimulation depends on the spontaneous activity within the neuronal network [31], we recorded the spontaneous neurotransmission at the baseline of our high-density neuronal culture by whole-cell patch-clamp technique. The resting membrane potential and other passive membrane properties such as capacitance and input resistance values were comparable to previous reports (**Figure EV 1I**) [32–35]. The neurons fired bursts of matured action potential spontaneously detected both in the current-clamp and voltage-clamp mode (**Figure 1D and 1E**) at day *in-vitro* 15. The action potential properties calculated through injecting a series of 500ms depolarizing current steps (−40pA to +540pA) were comparable to previously published reports (**Figure EV 1H, 1J, and 1K**) [33,34,36,37]. The presence of robust mEPSCs recorded at the baseline supports the presence of spontaneous excitatory neurotransmission (**Figure 1F and 1G**) [35]. Since the recording experiments were done with different batches of cultures, we sought to verify our observations in figure 1B in the same batch of cultures used for recording experiments. As observed before, glutamate addition led to a significant dip in the neuronal ATP/(ATP+ADP) ratio and was significantly rescued upon anisomycin treatment (**Figure EV 1M**) arguing that the observations presented in figure 1B and 1C were not the culture batch-specific artifacts.

In these neuronal cultures, we further measured what percentage of energy is utilized for endocytosis and ionic rebalancing as these are the two major mechanisms proposed to present the bulk of the energy burden within the cell [5,9]. We used TTX (1μM) to block the voltage-gated Na^+^ channels and small molecule inhibitors such as Ouabain (1mM) and Dynasore (100μM) to understand the contribution of the Na^+^/K^+^ ATPase activity and endocytosis respectively. Surprisingly, Dynasore treatment did not have a significant impact on glutamate-mediated energy usage (**Figure EV 1B and 1D**) while Ouabain and TTX reduced the energy consumption only marginally (**Figure EV 1C and 1E**). These results argued that the energy burden of protein synthesis outweighed that of the other mechanisms following neuronal activity.

We further examined the impact of mGluR and NMDAR stimulation individually on the post-synaptic ATP content considering that distinct sets of factors regulate local and global energy homeostasis within neurons [9]. For this, we prepared synaptoneurosomes from 30 days old (P30) rat cortices and measured synaptic ATP content on both the stimulations (**Figure 1H**). The ATP content was normalized to the total protein content, in this case, to account for the variability between samples. The significant enrichment of both pre-synaptic Synapsin 1 and post-synaptic PSD 95 proteins validated the preparation (**Figure 1I**). The synaptoneurosomes were also enriched with ‘snowman’ shaped pre and post-synaptic conglomerate (**Figure EV 1F**) [38] without a significant enrichment of glial protein GFAP (**Figure 1I**). In synaptoneurosomes, DHPG (100µM) treatment for 5 minutes led to a significant reduction in the ATP level. MPEP and anisomycin pre-incubation, however, diminished the effect of DHPG addition (**Figure 1J and Figure EV 1G**) as observed in Figure 1B and 1C. 5 minutes of NMDA (50µM) stimulation contrarily, had no significant impact on synaptic energy content (**Figure 1K and Figure EV 1G**). Therefore, mGluR dependent reduction in the neuronal ATP level and the ability of anisomycin to rescue the dip argue that protein synthesis shares the bulk of the glutamate-mediated energy burden.

### mGluR and NMDAR Impact Synaptic-dendritic ATP Levels with Distinct Kinetics

Since the glutamate receptors are concentrated on the dendritic spines [39], we hypothesized that they may not only influence the rate of global protein synthesis dynamically but may do so in a spatially distinct manner. We, therefore, monitored the changes in the ATP/ADP ratio live using ratiometric sensor PercevalHR [40] until 15 minutes after DHPG and NMDA addition both at the soma and at the distal dendrites (≥50µm away from the soma) of cortical neurons plated at low-density (**Figure 2A-2C**). Perceval pH bias was approximately corrected by simultaneous measurement of intracellular pH and establishing a linear relation between Perceval and pH-Red fluorescence as described previously [40,41] (**Figure EV 2A**). The low-density cultured neurons had a reduced level of spontaneous neurotransmission due to the lesser number of connections they formed [42–44] and showed reduced spiking frequency with increasing current steps injected (**Figure EV 1L**). However, less crowding within this kind of cell culture dishes allowed precise quantification of ATP/ADP ratio within dendritic compartments using microscopy-based techniques. We observed DHPG (50μM) application produced a significant and sustained drop in the dendritic ATP/ADP ratio within 2 minutes while NMDA (20μM) application produced a more delayed drop in the ratio after almost 10 minutes (**Figure 2C, 2E, and Figure EV 2B-2E**). To test the effect of protein synthesis inhibitors, we repeated both mGluR and NMDAR stimulation in the presence of anisomycin. Not only did anisomycin preincubation increase the baseline ATP level with time (**Figure EV 2F and 2G**), it significantly rescued the stimulation-mediated dip in the ATP level within the dendrites following the addition of both the stimuli (**Figure 2E and Figure EV 2C-2E**). These observations suggest that inhibition of protein synthesis by anisomycin increases the net ATP/ADP ratio of the cell. This gain in the ATP, however, can offset the relatively smaller extent of ATP consumption by mechanisms other than protein synthesis following glutamate receptor stimulation. To investigate the effect of anisomycin treatment on the surface expression of glutamate receptors, we quantified the cell surface level of mGluR5 and NR1 subunit of NMDAR in anisomycin treated cortical neurons. The surface mGluR5 level remained unchanged on anisomycin treated cells compared to basal untreated cultures while the NR1 surface level was up-regulated on anisomycin treatment (**Figure EV 1L and 1M**). These observations suggest that anisomycin pre-treatment does not reduce the surface level of glutamate receptors and that the observed recovery of the ATP/ADP ratio on both stimulations under anisomycin treated conditions was due to the inhibition of protein synthesis. Surprisingly both the drugs had little or no impact within the soma (**Figure 2B and 2D and Figure EV 2B**) indicating a more dynamic energy utilization within dendritic compartments compared to a more stable energy regulation at the soma [45].

**Figure 2.**
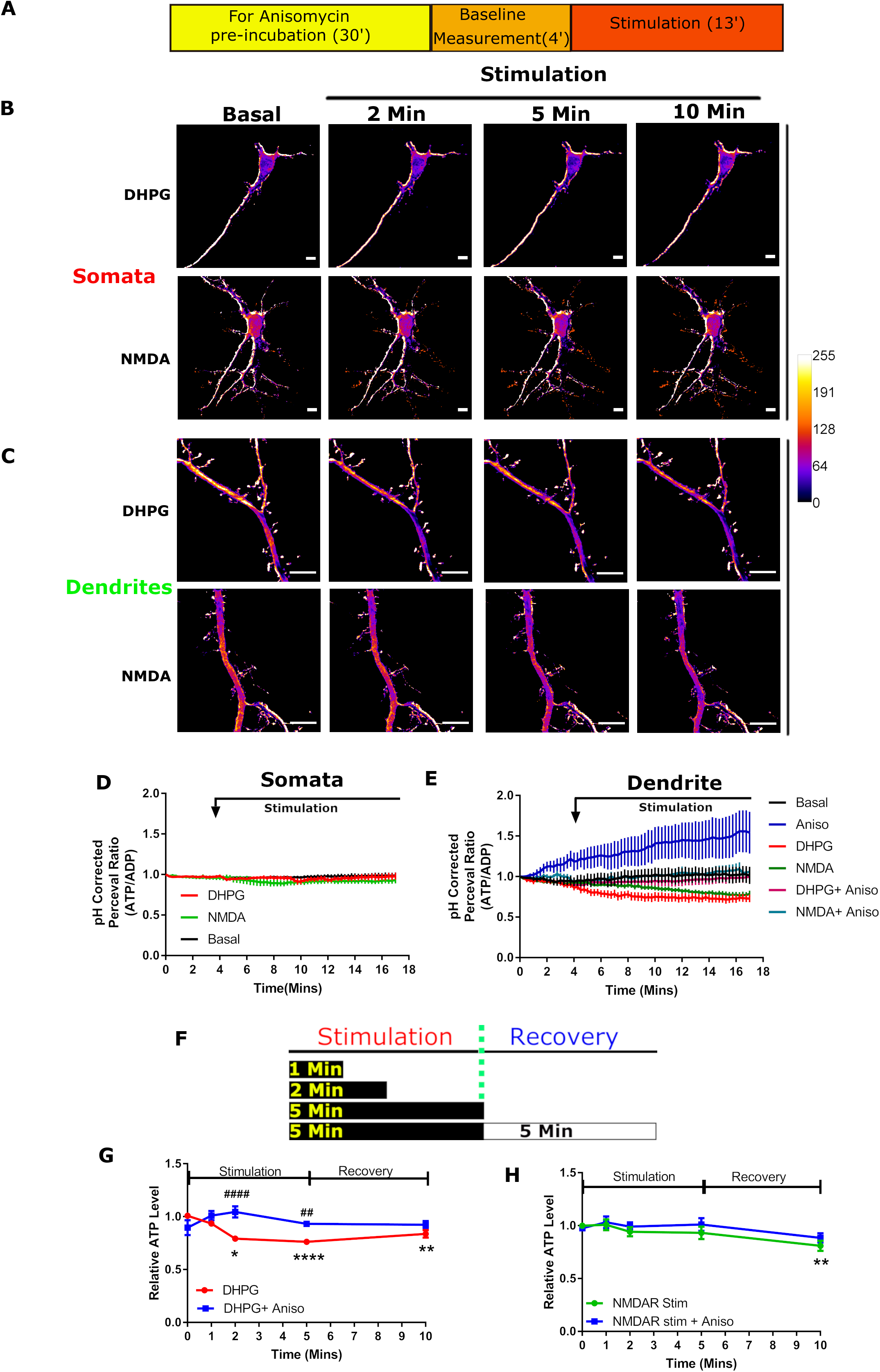
mGluR and NMDAR impact synaptic-dendritic ATP/ADP ratio with distinct Kinetics. **(A)** Schematic depicting the experimental workflow for the measurement of the ATP/ADP ratio using an imaging-based approach either from somatic or from dendritic compartments of cortical neurons (DIV 15) plated at low-density. **(B)** Representative neurons showing changes in the PercevalHR fluorescence ratio (∼ATP/ADP ratio) in somatic compartments on the bath application of DHPG and NMDA. Scale bar 10μm. **(C)** Representative neurons showing changes in the PercevalHR fluorescence ratio (∼ATP/ADP ratio) in dendritic compartments on the bath application of DHPG and NMDA. Scale bar 10μm. **(D)** Average traces depicting the time course of the normalized somatic ATP/ADP ratio on the basal conditions, on DHPG treatment, and on NMDA treatment. The curved arrow on the top indicates the time when the agonists were added and kept in the imaging media. Data presented as mean +/- SEM. Values in all other groups were normalized to the 0 min group for the corresponding experiment and all the data points are represented as a fraction of the initial time point. n= 5-6 cells from ≥3 independent platings. **(E)** Average traces depicting the time course of normalized dendritic ATP/ADP ratio on the basal condition, on the basal condition in the presence of anisomycin, on DHPG treatment, on NMDA treatment, on DHPG treatment in the presence of anisomycin and on NMDA treatment in the presence of anisomycin. The curved arrow on the top indicates the time when the agonists were added and kept in the imaging media. Data presented as mean +/- SEM. Values in all other groups were normalized to the 0 min group for the corresponding experiment and all the data points are represented as a fraction of the initial time point. n= 6-9 cells from ≥3 independent platings. **(F)** Schematic depicting stimulation protocol in cortical synaptoneurosomes for quantification of the synaptic ATP level. **(G)** Line graph showing the average normalized synaptic ATP level at various time points after DHPG treatment until 5 minutes of recovery in the presence or absence of anisomycin. Data presented as mean +/- SEM, *p<0.05, **p<0.01, ****p<0.0001 for comparison with 0 minute. ##p<0.01 and ####p<0.0001 for comparison between DHPG + anisomycin. Values in all other groups were normalized to the 0 min group for the corresponding experiment and all the data points are represented as a fraction of the initial time point. n≥ 5 animals per group. Two-way ANOVA followed by Bonferroni’s multiple comparison tests. **(H)** Line graph showing the average normalized synaptic ATP level at various time points after NMDA treatment until 5 minutes after recovery in the presence or absence of anisomycin. Data presented as mean +/- SEM, **p<0.01 for comparison with 0 minutes. Values in all other groups were normalized to the 0 min group for the corresponding experiment and all the data points are represented as a fraction of the initial time point. n≥ 4 animals per group. Two-way ANOVA followed by Bonferroni’s multiple comparison tests.

Since the steady-state ATP level is dependent on both the rate of production and consumption, we tested whether glutamate stimulation perturbs the ATP production and therefore leads to the observed drop in the neuronal ATP/ADP ratio. To verify, we quantified the dendritic ATP/ADP ratio on glutamate stimulation in the presence or absence of 2-deoxy glucose (2-DG, 30mM), a reversible inhibitor of glycolysis. In neurons, glycolysis supports the baseline ATP level [9] and thus 2-DG preincubation led to a significant decline in the basal dendritic ATP/ADP ratio (**Figure EV 2I and 2J**). However, 2-DG treatment did not alter the glutamate-mediated energy reduction in the dendritic ATP/ADP ratio (**Figure EV 2K and 2L**) suggesting the observed effect of glutamate stimulation is primarily because of ATP utilization and not because of altered ATP synthesis.

Further, we verified the effect of mGluR and NMDAR stimulation on the kinetics of ATP regulation at the mature synaptic compartments. We stimulated cortical synaptoneurosomes for a diverse period with DHPG and NMDA (1 min, 2 min, 5 min and allowed recovery for 5 minutes post-stimulation) and quantified the ATP levels (**Figure 2F and 1F**). As observed before, DHPG (100μM) addition rapidly decreased the synaptic ATP level within 2 minutes, which returned to the baseline by 5 minutes of recovery. Stimulation in the presence of anisomycin, however, failed to produce any significant change in the synaptic ATP level (**Figure 2G**) suggesting a dynamic alteration in protein synthesis creates a correlated change in the ATP level. NMDA stimulation had no significant impact on the ATP level until 5 minutes, as observed in dendrites (**Figure 2H, 2E and Figure EV 2H**). The effect of anisomycin preincubation had a comparable effect to that of NMDA treatment alone (**Figure 2H and Figure EV 2H**) suggesting an absence of active protein synthesis immediately following NMDAR stimulation. Together our observations establish that the well-correlated change in neuronal protein synthesis and ATP content follows distinct kinetics upon mGluR and NMDAR stimulation.

### mGluR and NMDAR Regulate eEF2 Phosphorylation to Create the Distinct Kinetics of Protein Synthesis

Since both NMDAR and mGluR impacted synaptic and dendritic ATP content dynamically in a protein synthesis-dependent manner, we decided to study the kinetics of *de-novo* protein synthesis following the addition of their specific agonists. We quantified the amount of newly-synthesized proteins using FUNCAT metabolic labeling based approaches as described previously [46]. Cortical neurons (DIV 15) plated at low-density were stimulated with DHPG or NMDA for metabolic labeling and the extent of labeling was quantified using mean fluorescence intensity which proportionally correlated with the rate of protein synthesis throughout the labeling period (**Figure 3A and 3B**). The absence of the FUNCAT signal in the control (without AHA) verifies that the signal is specific to AHA labeled new proteins (**Figure EV 3A**). We found that the DHPG application led to a significant elevation in the FUNCAT intensity which sustained till 5 min of recovery compared to the time-matched unstimulated control cultures (**Figure 3C, 3D, Figure EV 3A and 3B**). This demonstrated that mGluR stimulation activates robust protein synthesis in cortical neurons. In contrast, NMDA application caused a precipitous drop in the FUNCAT intensity within 2 minutes compared to the time-matched unstimulated control cultures. (**Figure 3C, 3D, Figure EV 3A and 3B**). We speculated that the observed reduction in the FUNCAT intensity was because of protein degradation [47][48][49]. To investigate this, we repeated NMDA stimulation for 2 minutes in the presence of MG132 (1µM), a 26s proteasome inhibitor that significantly improved the FUNCAT intensity compared to cultures treated with NMDA alone. This supported our hypothesis that a large-scale protein degradation ensues immediately following NMDAR stimulation. NMDA treatment for 20 minutes, however, led to an abundant increase in protein synthesis (**Figure EV 3C and 3D**) suggesting a dynamic modulation of global protein synthesis.

**Figure 3:**
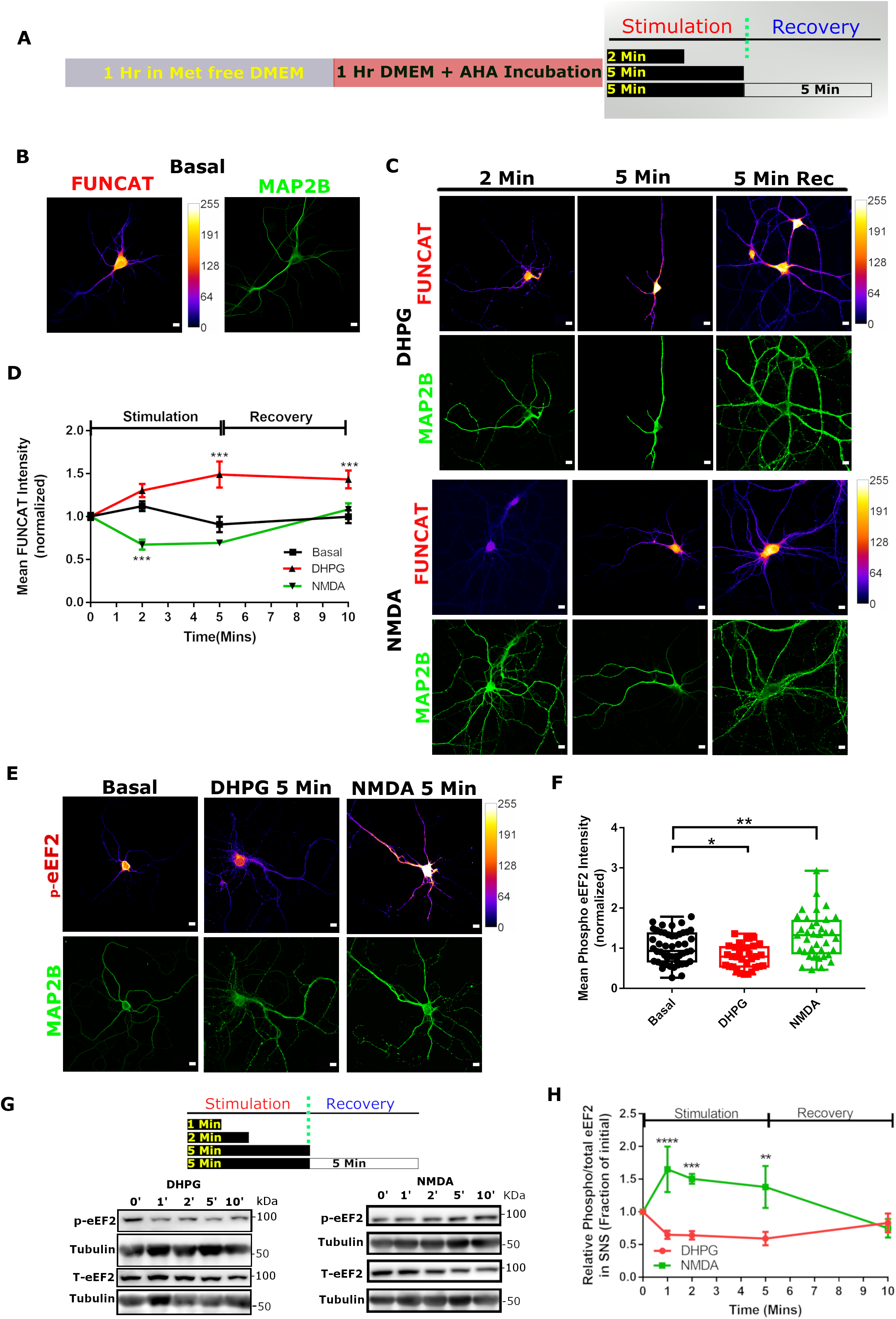
mGluR and NMDAR alter global translation by regulating eEF2 phosphorylation with distinct kinetics. **(A)** Schematic depicting the experimental workflow and the stimulation protocol to visualize and quantify newly synthesized proteins through metabolic labeling at various time points following DHPG or NMDA treatment in cortical neurons DIV 15 plated at low-density. **(B)** Representative images showing newly synthesized proteins on the basal condition visualized through FUNCAT metabolic labeling (pseudo-colored) in cortical neurons. MAP2B immunolabeling was used for identifying neurons and intensity was used for normalization. Scale bar 10μm. **(C)** Representative images showing newly synthesized proteins visualized through FUNCAT metabolic labeling (pseudo-colored) in cortical neurons at various time points following DHPG and NMDA treatment. MAP2B immunolabeling was used for identifying neurons and intensity was used for normalization. Scale bar 10μm. **(D)** Line graph showing the change in the average normalized FUNCAT intensity representing the quantity of newly synthesized proteins at various time points following DHPG and NMDA treatments. Data presented as mean +/- SEM. Data points in all groups were normalized to the average of the basal group. ***p<0.001 n= 21-54 neurons per group from 3 independent platings. Two-way ANOVA followed by Bonferroni’s multiple comparison test. **(E)** Representative images showing phospho-eEF2 immunolabeling (Pseudo-colored) in low-density cortical neurons on the basal condition, on DHPG treatment for 5 minutes and on NMDA treatment for 5 minutes. MAP2B immunolabeling was used for identifying neurons and intensity was used for normalization. Scale bar 10μm. **(F)** Box plot showing phospho-eEF2 normalized intensity distribution across multiple neurons on the basal condition, on DHPG treatment for 5 minutes and on NMDA treatment for 5 minutes. Data points in all groups were normalized to the average of the basal group. The box extends from 25^th^ to 75^th^ percentile with the middlemost line representing the median of the dataset. Whiskers range from minimum to maximum data point. *p<0.05, **p<0.01, n=31-49 cells per group from 3 independent platings. One-way ANOVA followed by Bonferroni’s multiple comparison test. **(G)** Representative immunoblots describing changes in the phospho-eEF2 and total-eEF2 levels at various time points after DHPG and NMDA treatment in cortical synaptoneurosomes. Note in each case phospho and total eEF2 levels were normalized individually to their respective tubulin levels for calculating the phospho/total eEF2 ratio. **(H)** Line graph showing the average value of the synaptic phospho/total ratio of eEF2 at various time points after DHPG and NMDA treatment until 5 minutes of recovery. Data presented as mean +/- SEM, **p<0.01. Values in all other groups were normalized to the 0 min group for the corresponding experiment and all the data points are represented as a fraction of the initial time point. ***p<0.001, ****p<0.0001, n≥ 5 animals per group. Two-way ANOVA followed by Bonferroni’s multiple comparison test.

Since translation can be regulated at multiple stages, we decided to focus on the elongation regulation as elongation block is a viable mechanism to modulate translation within a cell [50] and plays an important role in the context of NMDAR mediated protein synthesis [21,51]. The rate of ribosomal translocation can be regulated by altering the phosphorylation of eEF2 [52]. Hyperphosphorylation eEF2 at Thr^56^ mediated by eEF2K reduces the rate of translation elongation [53–55]. Therefore, at any given instance, the status of eEF2 phosphorylation reflects an integrated response from multiple biochemical pathways [21,56] and prompted us to investigate the activity-dependent modulation of eEF2 phosphorylation through immunolabeling 5 minutes after both mGluR or NMDAR stimulation in cortical neurons (**Figure 3E**). We observed a significant reduction in the p-eEF2 immunolabeling 5 minutes after DHPG treatment and a significant elevation in the immunolabeling 5 minutes after NMDA treatment compared to the time-matched unstimulated control cultures (**Figure 3F**). This indicated that eEF2 phosphorylation was tuned to create receptor-specific protein synthesis response and that at any given instance eEF2 phosphorylation reliably reflected the status of protein synthesis within a cell. Hence, we used the phospho/total ratio of eEF2 as a readout for global translation in cortical synaptoneurosomes (**Figure 3G and 3H**) and high-density cortical neurons (**Figure EV 3E and 3F**). Both synaptoneurosomes and neurons were stimulated for different periods as mentioned before (**Figure 2F**). We observed that a significant difference exists between the kinetics of how eEF2 phosphorylation was altered upon mGluR and NMDAR stimulation (**Figure 3G and Figure EV 3F**). Comparison with the basal condition revealed that mGluR stimulation led to an immediate and sustained reduction in the phospho/total eEF2 ratio while NMDAR stimulation led to an immediate increase in the ratio both in the synaptoneurosomes and in cultured neurons. The phosphorylation status returned to the baseline by 5 mins of recovery in both the cases (**Figure 3H and Figure EV 3E, 3F and 4A**). The temporally-matched inverse correlation between FUNCAT signal and eEF2 phosphorylation implied that the glutamate receptors had a strong influence over eEF2 to modulate protein synthesis dynamically.

### Activity-dependent Dynamic Modulation of AMPK Is Necessary to Alter the eEF2 Phosphorylation

The phosphorylation of eEF2 is regulated by eEF2 Kinase which is known to be a substrate for AMP-activated protein kinase [57]. AMPK is reported to sense the intracellular AMP/ATP ratio or the ADP/ATP ratio and inhibits protein synthesis during energy stress [28]. Therefore, we sought to understand if the glutamate receptors needed to alter AMPK function to regulate protein synthesis. Since phosphorylation of Thr^172^ is known to directly correlate with AMPK activation [58], we examined the phosphorylation status of AMPK through immunolabeling 5 minutes after mGluR and NMDAR stimulation in cortical neurons (**Figure 4A**). We observed a significant reduction in the p-AMPK immunolabeling upon mGluR stimulation while an elevation in the p-AMPK level following NMDAR stimulation compared to the time-matched unstimulated control cultures (**Figure 4B**). This signified that the glutamate receptors held the ability to modulate AMPK function following their activation. To verify whether mGluR and NMDAR alter AMPK phosphorylation dynamically, we quantified the phospho/total ratio of AMPK from cortical synaptoneurosomes after various periods of DHPG or NMDA treatment (**Figure 4C**). We observed a persistent and significant reduction in the phospho/total ratio of AMPK within 2 minutes of DHPG application compared to unstimulated synaptoneurosomes (**Figure 4D and Figure EV 4A**). Surprisingly, DHPG mediated reduction in AMPK phosphorylation was reversed in the presence of anisomycin (**Figure 4C, 4D and Figure EV 4B**) indicating that the AMPK activity is regulated in turn by the newly synthesized proteins involving a feedback-inhibition. We speculated, therefore, that the synthesis of any AMPK specific phosphatase could explain the dephosphorylation of AMPK.

**Figure 4:**
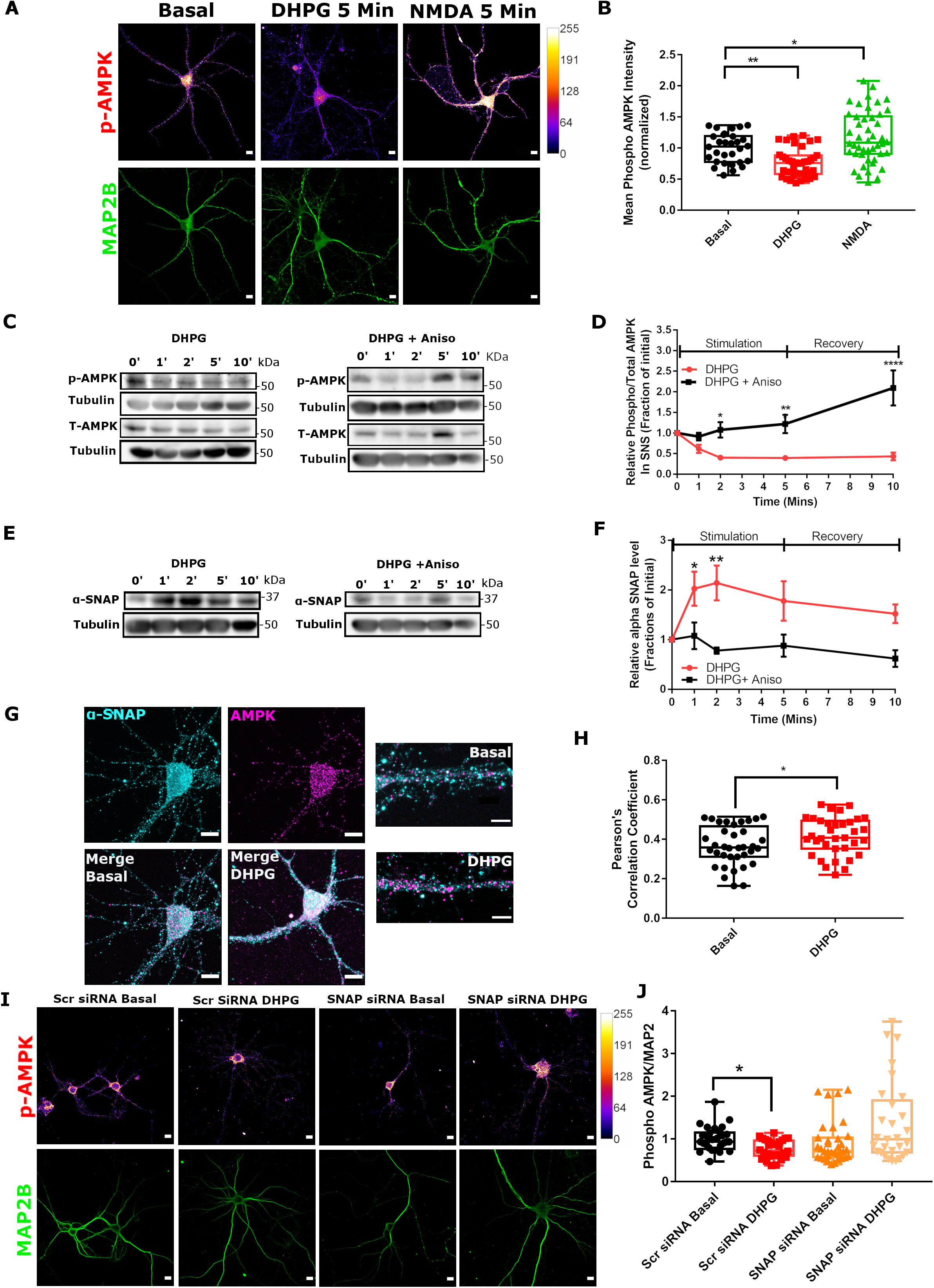
mGluR modulates AMPK function through a protein synthesis-dependent feedback mechanism: **(A)** Representative images showing phospho-AMPK immunolabeling (Pseudo-colored) in low-density cortical neurons on the basal condition, on DHPG treatment for 5 minutes and on NMDA treatment for 5 minutes. MAP2B immunolabeling was used for identifying neurons and intensity was used for normalization. Scale bar 10μm. **(B)** Box plot showing the normalized phospho-AMPK intensity distribution across multiple neurons on the basal condition, on DHPG treatment for 5 minutes and on NMDA treatment for 5 minutes. Data points in all groups were normalized to the average of the basal group. The box extends from 25^th^ to 75^th^ percentile with the middlemost line representing the median of the dataset. Whiskers range from minimum to maximum data point. *p<0.05, **p<0.01, n=30-46 cells per group from 3 independent platings. One-way ANOVA followed by Bonferroni’s multiple comparison test. **(C)** Representative immunoblots describing changes in the phospho-AMPK and total-AMPK levels at various time points after DHPG treatment in cortical synaptoneurosomes in the presence or absence of anisomycin. Note in each case phospho and total AMPK levels were normalized individually to their respective tubulin levels before calculating the phospho/total ratio of AMPK. **(D)** Line graph showing the normalized average value of the synaptic phospho/total ratio of AMPK at various time points after DHPG treatment in the presence or absence of anisomycin. Data presented as mean +/- SEM. Values in all other groups were normalized to the 0 min group for the corresponding experiment and all the data points are represented as a fraction of the initial time point. *p<0.05, **p<0.01, ****p<0.0001, n= 5 animals per group. Two-way ANOVA followed by Bonferroni’s multiple comparison test. **(E)** Representative immunoblots depicting the changes in the α-SNAP protein levels at various time points after DHPG treatment in cortical synaptoneurosomes in the presence or absence of anisomycin. **(F)** Line graph showing the normalized average value of the synaptic α-SNAP protein levels at various time points after DHPG treatment in the presence or absence of anisomycin. Data presented as mean +/- SEM. *p<0.05. Values in all other groups were normalized to the 0 min group for the corresponding experiment and all the data points are represented as a fraction of the initial time point. **p<0.01, n≥ 4 animals per group. Two-way ANOVA followed by Bonferroni’s multiple comparison test. **(G)** Representative images showing the α-SNAP (cyan) and AMPK (magenta) immunolabeling in low-density cortical neurons. Merge shows the colocalization of the two channels both on the basal condition and on DHPG treatment for 5 minutes. Scale bar 10μm. Zoomed in representative images of dendrites showing the merge of both channels on the basal and DHPG treated conditions. Scale bar 5μm. **(H)** Box plot depicting the quantification of co-localization through Pearson’s correlation coefficient between α-SNAP and AMPK in cortical neurons on the basal condition and on DHPG treatment for 5 minutes. The box extends from 25^th^ to 75^th^ percentile with the middlemost line representing the median of the dataset. Whiskers range from minimum to maximum data point. *p<0.01, n≥35 cells per group from 4 independent platings. Unpaired-sample t-test. **(I)** Representative images showing the phospho-AMPK immunolabeling (Pseudo-colored) in low-density cortical neurons on the basal condition and on DHPG treatment for 5 minutes in the presence of scrambled siRNA and in the presence of α-SNAP siRNA. MAP2B immunolabeling was used for identifying neurons and intensity was used for normalization. Scale bar 10μm. **(J)** Box plot showing the normalized phospho-AMPK intensity distribution across multiple neurons on the basal condition in the presence of scrambled siRNA, on DHPG treatment for 5 minutes in the presence of scrambled siRNA, on the basal condition in the presence of α-SNAP siRNA, on DHPG treatment for 5 minutes in the presence of α-SNAP siRNA. Data points in all groups were normalized to the average of the scrambled siRNA basal group. The box extends from 25^th^ to 75^th^ percentile with the middlemost line representing the median of the dataset. Whiskers range from minimum to maximum data point. *p<0.05, n=29-35 cells per group from 3 independent platings. Kruskal-Wallis test followed by Dunn’s multiple comparison test.

While the exact identity of the AMPK phosphatase is still elusive [59–61], we chose to investigate the role of α-SNAP, an AMPK specific inhibitor [61], in the context of mGluR stimulation in cortical synaptoneurosomes (**Figure 4E**). We observed a significant elevation in the α-SNAP level within 1 minute of DHPG addition, which was absent in the anisomycin treated preparations (**Figure 4F**). The activity-induced changes in α-SNAP level inversely correlated with the kinetics of AMPK activation (**Figure 4D**) suggesting that α-SNAP plays a critical role in dictating the status of AMPK activity upon mGluR stimulation. We sought to confirm the role of α-SNAP with two approaches. First, we quantified the colocalization between α-SNAP and AMPK, which increased modest yet significantly on mGluR stimulation in low-density cultured neurons (**Figure 4G and 4H**). Second, we acutely knocked down the α-SNAP protein level using the siRNA-based approach and quantified the p-AMPK levels in the neurons (**Figure 4I**). The α-SNAP siRNA treatment did not alter the p-AMPK level significantly on the basal condition while it led to a marked reduction in the α-SNAP protein level compared to scrambled siRNA treated cultures (**Figure EV 4C**). However, α-SNAP-siRNA treatment eliminated the DHPG induced reduction in the AMPK phosphorylation (**Figure 4J**) indicating a more complex regulation of AMPK. Our observations also argue that the recruitment of α-SNAP for the AMPK regulation is an exclusive feature of the mGluR mediated signal transduction.

### NMDAR Regulates AMPK Activity in a Ca2+ Dependent Manner

The fact that NMDAR stimulation led to an up-regulation of the p-AMPK level (**Figures 4A and 4B**) and that AMPK is a known substrate of CamKKIIβ [62], made us wonder whether the entry of extracellular Ca^2+^ through open NMDAR channels regulates AMPK phosphorylation. We first decided to investigate the status of the AMPK phosphorylation following various periods of NMDA incubation (**Figure 5A**). We observed a rapid and persistent increase in the phospho/total ratio of AMPK within 1 minute of NMDA addition (**Figure 5B**) corroborating our previous observation in Figure 4B. We confirmed the role of Ca^2+^ by repeating the stimulation in the absence of extracellular Ca^2+^, which eliminated the NMDAR dependent elevation of the phospho/total ratio of AMPK (**Figure 5B**). Besides, the NMDAR mediated increase in the phospho/total ratio of eEF2 was diminished significantly in the absence of extracellular Ca^2+^ indicating AMPK and eEF2 phosphorylations were altered in a correlated fashion on NMDAR stimulation with Ca^2+^ playing a critical role in dictating the kinetics (**Figure EV 4E and 4F**). We sought to verify the NMDA dependent Ca^2+^ entry by live monitoring the cytosolic free Ca^2+^ following NMDA addition in the low-density cultured neurons using the Fluo-8AM probe. We observed a rapid and persistent increase in Fluo8 fluorescence on NMDA addition (**Figure 5C**) indicating an increase in the cytosolic Ca^2+^ concentration on NMDA treatment. The rise in Fluo8 fluorescence could be reversed by D-AP5 or MK801 pre-incubation for 30 minutes (**Figure 5D, 5E, and Figure EV 4D**) arguing the response was NMDAR specific. A further increase in fluorescence on ionomycin addition in the presence of 10mM Ca^2+^ provided the fluorescence maxima for each cell (**Figure 5D**). The elevation in fluorescence was significantly reduced in the absence of extracellular Ca^2+^ (**Figure 5D, 5E, and Figure EV 4D**) indicating that a large amount of extracellular Ca^2+^ enters the cell upon NMDA addition. This elevation in cytosolic free Ca^2+^ activated AMPK in response to stimulation possibly by engaging CamKKIIβ [51,63]. Interestingly, we observed a similar extent of elevation in the phospho/total ratio of eEF2 on NMDAR stimulation in cortical synaptoneurosomes both in the presence or absence of extracellular Mg^2+^, an ion that occludes the NMDAR channel pore (**Figure EV 4G**). Together these results suggested that NMDA addition led to an abundant Ca^2+^ entry through the open channels of synaptic NMDAR, regulating both AMPK and eEF2 phosphorylation within the neuron.

**Figure 5:**
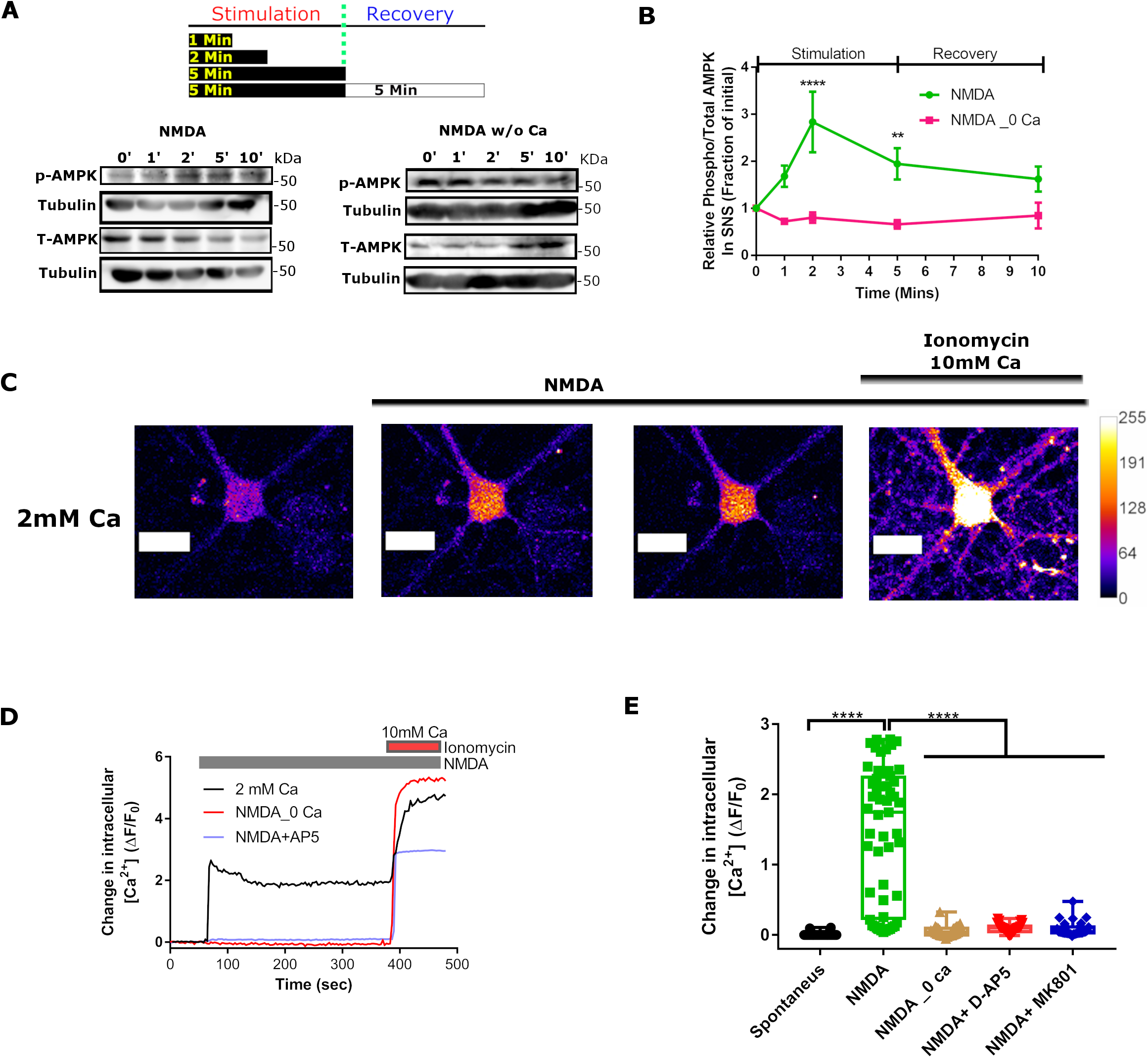
NMDAR mediated AMPK activation is extracellular Ca^2+^-dependent. **(A)** Representative immunoblots describing changes in the phospho-AMPK and total-AMPK levels at various time points after NMDA treatment in cortical synaptoneurosomes in the presence or absence of extracellular Ca^2+^. Note in each case phospho and total AMPK levels were normalized individually to their respective tubulin levels before calculating the phospho/total AMPK ratio. **(B)** Line graph showing the normalized average value for the synaptic phospho/total ratio of AMPK at various time points after NMDA treatment until 5 minutes of recovery in the presence or absence of extracellular Ca^2+^. Data presented as mean +/- SEM. Values in all other groups were normalized to the 0 min group for the corresponding experiment and all the data points are represented as a fraction of the initial time point. **p<0.01, ****p<0.0001, n= 6 animals per group. Two-way ANOVA followed by Bonferroni’s multiple comparison test. **(C)** Representative images depicting intracellular Ca^2+^ levels through Fluo8 fluorescence (Pseudo-colored) of a cortical neuron plated at low-density before stimulation, 15 sec (immediate) after NMDA treatment, 300 sec (delayed) after NMDA treatment and after ionomycin treatment in the presence of 10mM Ca^2+^. Scale bar 20μm. **(D)** Representative time trace showing the normalized change in Fluo8 fluorescence on NMDA treatment in the presence or absence of extracellular Ca^2+^ and D-AP5. Values in all other groups were normalized to the unsimulated condition for the corresponding experiment and all the data points are represented as a fraction of the initial time point. Ionomycin treatment in the presence of 10mM Ca^2+^ was used to calculate fluorescence maximum for a particular cell. **(E)** Box plot showing the distribution of ΔF/F_0_ across multiple neurons on the basal condition, on NMDA treatment in the presence Ca^2+^, on NMDA treatment in the absence of Ca^2+^, on NMDA treatment with D-AP5 and on NMDA treatment along with MK-801. The box extends from 25^th^ to 75^th^ percentile with the middlemost line representing the median of the dataset. Whiskers range from minimum to maximum data point. ****p<0.0001, n≥30 neurons per group from 3 independent platings. One-way ANOVA followed by Bonferroni’s multiple comparison test.

### Acute Perturbation of AMPK Disrupts the Receptor-specific Translation Response

The link between the AMPK regulation and glutamate receptor-mediated protein synthesis in cortical neurons prompted us to investigate whether the AMPK regulation was necessary to generate the receptor-specific protein synthesis response. We probed this question using two independent approaches. To begin with, we repeated the mGluR stimulation in the presence of 5-aminoimidazole-4-carboxamide ribonucleotide (AICAR; 1mM) which is known to activate AMPK acutely [64] (**Figure 6A**). We observed AICAR pre-treatment for 1 hour led to a significant elevation in p-eEF2 levels (**Figure 6C, Figure EV 4H**) without affecting the FUNCAT signal (**Figure 6D, Figure EV 4H**) basally in the low-density cortical neurons. However, AICAR treatment significantly rescued the DHPG induced reduction in p-eEF2 levels (**Figure 6A, B, Figure EV 4H**) and eliminated the DHPG induced rise in the FUNCAT intensity (**Figure 6C, D, Figure EV 4H**). This meant that the acute apriori activation of AMPK disrupts the mGluR-mediated dephosphorylation of eEF2 and a subsequent elevation in protein synthesis. Similarly, we examined the impact of AMPK activation on NMDAR mediated protein synthesis regulation. We stimulated cortical neurons with NMDA in the presence of compound C (Dorsomorphin/CC), a small molecule inhibitor of AMPK (**Figure 6E**). CC pre-treatment for 1 hour had no significant impact on p-eEF2 levels (6F, Figure EV 4H) but led to a significant increase in the FUNCAT intensity (6G, Figure EV 4H). However, CC abolished the NMDA mediated rise in p-eEF2 levels (**Figure 6F**) and significantly rescued the inhibition of protein synthesis (**Figure 6G**). Therefore, acute inhibition of AMPK perturbed the NMDA mediated hyperphosphorylation of eEF2 and inhibition of global protein synthesis. These also confirmed the pivotal role of AMPK in dictating the fate of global protein synthesis following glutamate receptor activation.

**Figure 6:**
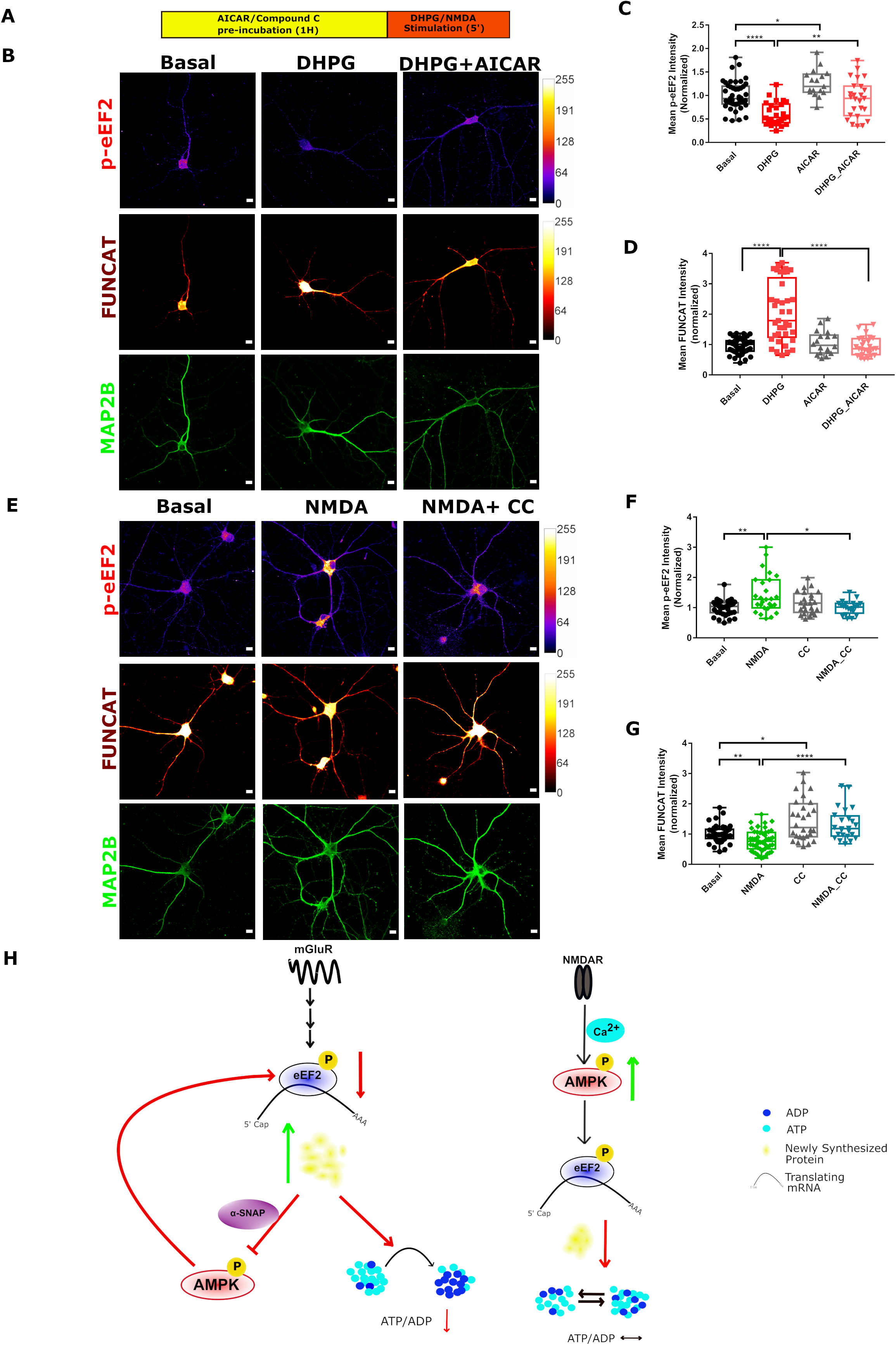
Perturbation of AMPK function disrupts receptor-specific translation response: **(A)** Schematic depicting the experimental workflow for quantifying p-eEF2 levels and FUNCAT intensity changes on DHPG and NMDA treatment in low-density cortical neurons (DIV 15). **(B)** Representative images showing the phospho-eEF2 immunolabeling (Pseudo-colored) and newly synthesized proteins as FUNCAT signal (pseudo-colored) in cortical neurons on the basal condition and on DHPG (50μM) treatment for 5 minutes in the presence or absence of AICAR (1mM). MAP2B immunolabeling was used for identifying neurons and intensity was used for normalization. Scale bar 10μm. **(C)** Box plot showing the normalized phospho-eEF2 intensity distribution across multiple neurons on the basal condition, on DHPG treatment for 5 minutes, on AICAR treatment and on DHPG treatment with AICAR. Data points in all groups were normalized to the average of the basal group. The box extends from 25^th^ to 75^th^ percentile with the middlemost line representing the median of the dataset. Whiskers range from minimum to maximum data point. *p<0.05, **p<0.01, ****p<0.0001, n=17-46 cells per group from 3 independent platings. One-way ANOVA followed by Bonferroni’s multiple comparison test. **(D)** Box plot showing the FUNCAT intensity distribution across multiple neurons on the basal condition, on DHPG treatment for 5 minutes, on AICAR treatment and on DHPG treatment with AICAR. Data points in all groups were normalized to the average of the basal group. The box extends from 25^th^ to 75^th^ percentile with the middlemost line representing the median of the dataset. Whiskers range from minimum to maximum data point. ****p<0.0001, n=18-57 cells per group from 3 independent platings. Kruskal-Wallis test followed by Dunn’s multiple comparison test. **(E)** Representative images showing the phospho-eEF2 immunolabeling (Pseudo-colored) and newly synthesized proteins as FUNCAT signal (Pseudo-colored) in cortical neurons on the basal condition and on NMDA treatment for 5 minutes in the presence or absence of Compound C. MAP2B immunolabeling was used for identifying neurons and intensity was used for normalization. Scale bar 10μm. **(F)** Box plot showing phospho-eEF2 intensity distribution across multiple neurons on basal condition (Vehicle control), on NMDA treatment for 5 minutes, on Compound C treatment and on NMDA treatment with Compound C. Data points in all groups were normalized to the average of the basal group. The box extends from 25^th^ to 75^th^ percentile with the middlemost line representing the median of the dataset. Whiskers range from minimum to maximum data point. *p<0.05, **p<0.01, n=21-36 cells per group from 3 independent platings. Kruskal-Wallis test followed by Dunn’s multiple comparison test. **(G)** Box plot showing the FUNCAT intensity distribution across multiple neurons on the basal condition, on NMDA treatment for 5 minutes, on Compound C treatment and on NMDA treatment with Compound C. Data points in all groups were normalized to the average of the basal group. The box extends from 25^th^ to 75^th^ percentile with the middlemost line representing the median of the dataset. Whiskers range from minimum to maximum data point. *p<0.05, **p<0.01, ****p<0.0001, n=24-52 cells per group from 3 independent platings. Kruskal-Wallis test followed by Dunn’s multiple comparison test. **(H)** Model of the receptor-specific regulation of AMPK-eEF2 signaling axis and their subsequent effect on global translation. mGluR stimulation (Left) inactivates AMPK, a necessity for the dephosphorylation of eEF2. This, in turn, leads to an enhanced protein synthesis thereby leading to the consumption of ATP. NMDAR stimulation (Right), however, led to an activation of AMPK by allowing the entry of extracellular Ca^2+^ through open NMDAR channels. This resulted in an increase in the eEF2 phosphorylation and an inhibition of global protein synthesis having no significant impact on neuronal energetics.

## Discussion

In our work, we demonstrate that the activity-mediated protein synthesis in response to mGluR and NMDAR stimulation leads to distinct and dynamic alterations of the neuronal energy level. A unique combination of AMPK-eEF2 signaling brings about the characteristic changes in protein synthesis specific for each stimulus.

### Protein Synthesis Consumes Bulk of the Energy on Glutamate Stimulation in Neurons

Protein synthesis is essential for axonal path finding [65], axonal and dendritic branching [66,67] synaptic plasticity [68,69] and other neuronal functions. Our study establishes protein synthesis to have a major contribution to activity-mediated energy consumption. However, it would be interesting to delineate whether any secondary or tertiary mechanism activated following protein synthesis is responsible for any fraction of this consumption. Strategies allowing the synthesis of non-functional proteins using genetically incorporated un-natural amino acids can assist in addressing this issue [70]. Our observation also suggests that the energetic cost Na+/K+ ATPase is less than protein synthesis unlike predicted earlier [5]. Though vesicle endocytosis is known to cause the major energy drainage at the presynaptic nerve terminals, we did not observe any significant contribution originating from it at the post-synaptic compartments [9]. Our observations, however, demand further exploration of the energy consumption by other activated synaptic mechanisms such as organellar movement, cytoskeletal rearrangement, autophagy, global protein degradation and others [5,71].

### mGluR and NMDAR Affect the Rate of Global Translation

Results from our work also reveal that mGluR dependent protein synthesis follows distinct kinetics as compared to that of NMDAR. The kinetics of protein synthesis was inversely correlated with the kinetics of ATP consumption. Group I mGluR have been widely reported to activate protein synthesis [19,20,72] across various brain regions [73–75]. Our results indicate that mGluR mediated translation activation produced a rapid reduction in the synaptic-dendritic ATP level. However, the somatic ATP level remained unchanged on mGluR stimulation even though protein synthesis was activated in the cell body (**Figure 3C and 3D**). This implies that the impact of protein synthesis on the steady-state ATP level depends on how rapidly the local energy supply pathways compensate for the enhanced energy demand within a neuronal compartment [8,76]. In contrast, NMDAR stimulation led to a biphasic protein synthesis response with an early inhibition and a delayed activation phase [21]. It is interesting that unlike the effect of anisomycin (**Figure EV 2F and 2G**), NMDAR mediated inhibition of protein synthesis did not elevate the cellular ATP level and only produced a delayed reduction of the same. This is probably because of the ATP consumed for large scale protein degradation [77] (**Figure EV 3D**) offsets the gradual build-up of ATP upon protein synthesis inhibition (**Figure EV 2G**).

### The role of AMPK-eEF2 Signaling to Create Differential Response to mGluR and NMDAR Stimulation

Our results also emphasize the key role of AMP-activated protein Kinase in both mGluR and NMDAR mediated translation regulation. Previously, AMPK has been demonstrated to regulate activity-induced energy metabolism [25], sustained activity-induced GLUT4 membrane expression in nerve terminals [78] and various forms of synaptic plasticity [29,30]. Our current observations, however, establish AMPK as a mechanistic link to coordinate activity-induced protein synthesis and energy metabolism. AMPK inhibits global protein synthesis by activating eEF2 Kinase directly or via indirect mechanisms [79]. mGluR mediated AMPK inhibition, therefore, allows the dephosphorylation of eEF2 causing an enhanced protein synthesis. We found that α-SNAP, one of the proteins synthesized on mGluR stimulation, inhibited AMPK likely through its phosphatase activity [61] suggesting there exist several noncanonical mechanisms to regulate AMPK within neurons. Also, it would be interesting to understand the contribution of other AMPK phosphatases like PP2A in governing the course of mGluR-induced signaling and protein synthesis [80]. NMDAR stimulation, on the contrary, led to immediate activation of AMPK in a Ca2+ dependent manner probably by recruiting the upstream CamKKIIβ. The activated AMPK, in turn, led to hyperphosphorylation of eEF2 (**Figure EV 4D**) resulting in an inhibition of neuronal protein synthesis as observed before [51].

In summary, our study demonstrates the striking influence of protein synthesis on the synaptic-dendritic energy homeostasis following the stimulation of glutamate receptors. AMPK plays a pivotal role in dictating this correlation (**Figure 6H**). It would be intriguing, however, to elucidate how the correlation is altered in neurodevelopment and various pathophysiological conditions such as stroke, epilepsy or neurodegenerative diseases. The study may also help in identifying potential therapeutic targets for diseases involving bioenergetic impairments and dysregulated protein synthesis.

## Materials and Methods

### Ethics Statement

All animal work was done in compliance with the procedures approved by the Institutional Animal Ethics committee (IAEC) and the Institutional Biosafety Committee (IBSC), InStem, Bangalore, India. All rodent work was done with Sprague Dawley (SD)rats. Rats were kept in 20-22’c temperature, 50-60 relative humidity, 0.3μm HEPA filtered air supply at 15-20 ACPH and 14/10 light/dark cycle maintained. Food and water were provided *ad libitum*.

### Antibodies, Drugs, and Other Reagents

Anti-phospho eEF2 antibody (Thr 56; cat no: 2331; Used at 1:1000 dilution for western blotting analysis and 1:250 for immunocytochemistry), anti-eEF2 antibody (cat. no 2332; Used at 1:1000 dilution for western blotting analysis), anti-phospho AMPKα antibody (Thr 172; cat. no: 2535; used at 1:500 dilution for western blotting analysis and 1:100 for immunocytochemistry analysis) and anti-AMPK antibody (cat no: 2532; used at 1:1000 dilution for western blotting analysis) were obtained from Cell Signaling Technologies (MA, US). Anti-tubulin antibody (cat no: T9026; used at 1:1000 dilution for western blotting analysis), anti-MAP2B antibody (cat no: M9942; used at 1:500 dilution for immunocytochemistry), anti-rabbit HRP labeled secondary antibody (cat no: A0545; used at 1:5000 dilution for western blotting analysis) and anti-mouse HRP labeled secondary antibody (cat no: A9044; used at 1:5000 dilution for western blotting analysis) were obtained from Sigma-Aldrich (St. Louis, MO). Anti-rabbit secondary antibody Alexa 555 labeled (cat no: A11032; used at 1:500 dilution for immunocytochemistry), anti-mouse secondary antibody Alexa 488 labeled (cat no: A11008; used at 1:500 dilution for immunocytochemistry) and α-SNAP (NAPA) siRNA were obtained from Thermo Fisher Scientific (Waltham, MA). Anti-α-SNAP antibody (cat no: X1026; used at 1:1000 dilution for western blotting analysis) was obtained from Exalpha Biologicals. Anti-Synapsin 1 antibody (cat no: ab64581, Used at 1:1000 dilution for western blotting) was obtained from Abcam (Cambridge, UK). Glutamate (25μM), NMDA (20μM for neurons and 40μM for SNS), CNQX (40μM), Anisomycin (25μM for neurons and 50μM for synaptoneurosomes), Cycloheximide (100μg/ml), Ouabain octahydrate (1mM), Dynasore hydrate (100μM), Poly-L-Lysine (0.2mg/ml), Pluronic-F-127(0.002%), BAPTA (10mM), Ionomycin (10μM), 2 deoxy-glucose (30mM) and oligomycin A (2.5μM) were obtained from Sigma-Aldrich (St. Louis, MO). S-3,5 DHPG (20μM for neurons and 100μM for SNS), 5-aminoimidazole-4-carboxamide ribonucleotide (AICAR, 1mM), D-AP5 (25μM), MK-801 (10μM), MPEP (10μM), Tetrodotoxin citrate [TTX, 1μM for measuring (ATP/ATP+ADP) and 0.5 μM for electrophysiology experiments] were obtained from Tocris Biosciences. Compound C (dorsomorphin; 40μM in synaptoneurosomes and 20μM in cultured neurons) were obtained from Santa Cruz Biotechnology (Santa Cruz, CA). Fluo-8 AM (2μM) was obtained from AAT Bioquest (Sunnyvale, CA). FUGW-PercevalHR was a gift from Gary Yellen (Addgene plasmid # 49083; http://n2t.net/addgene: 49083; RRID: Addgene_49083). GW1-pHRed was a gift from Gary Yellen (Addgene plasmid # 31473; http://n2t.net/addgene: 31473; RRID: Addgene_31473).

### Synaptoneurosome Preparation

Synaptoneurosomes were prepared from SD WT rat cortices at the age between post-natal day 28-33 (P28-33) following differential centrifugation method as described previously [21,81,82]. Briefly, rat cortices were dissected and homogenized at 4°C in 8 volumes of synaptoneurosome (SNS) homogenization buffer [containing (in mM) 25 Tris-HCl pH 7.4, 118 NaCl, 4.7 KCl, 1.2 MgSO4, 2.5 CaCl2, 1.53 KH2PO4, and 212.7 glucose, supplemented with Complete protease inhibitors (Roche)]. Homogenates were then passed through three 100μm nylon mesh filters followed by one 11μm filter MLCWP 047 Millipore (Bedford, MA) and centrifuged at 1000g for 20 minutes at 4°C. The pellet was resuspended into 2ml of the same buffer. In experiments with no Ca2+, CaCal2 was removed from the resuspension buffer. The resuspended SNS particles were then incubated at 37°C to regain active metabolism.

### Synaptoneurosome Stimulation

For Stimulation, synaptoneurosomes were pre-warmed at 37°C for a minimum of 5 minutes. A sample fraction aliquoted at this point was considered as the unstimulated condition or 0 min of stimulation for further analysis. This was followed by Gp I mGluR and NMDAR stimulation with specific agonists (s)3,5-dihydroxyphenylglycine (DHPG; 100μM) and N-methyl D-aspartate (NMDA; 40μM) respectively. Further fractions were collected after 2 minutes and 5 minutes of stimulation. A fraction was collected 5 minutes post-stimulation (5 min stimulation + 5 min recovery). For this, the synaptoneurosomes were first stimulated for 5 minutes with agonists. This is followed by the removal of the stimulus through centrifugation at 13,000g for 20 seconds and resuspension with 37°C pre-warmed synaptoneurosome homogenization buffer. These synaptoneurosomes were further incubated for 5 minutes and processed for ATP measurement, protein estimation, and western blot analysis. For ATP measurements, stimulation was terminated by direct lysis of synaptoneurosomes with an equal volume of boiling water [83]. This was followed by centrifugation at 20,000g for 2.5 minutes for the measurement of soluble ATP content from the supernatant. For protein estimation and western blot analysis, synaptoneurosomes fractions were centrifuged at 13,000g for 20 seconds followed by lysis with lysis buffer [containing: 50 Tris-Cl (pH-7.4), 150 NaCl, 5 MgCl2, 1% Triton-X-100, supplemented with EDTA free protease inhibitor complex (Sigma, cat. no. S8830) and phosphatase inhibitor cocktail (Roche, ref.no. 04906837001)] (Stimulation protocol figure 2F). In experiments involving pre-treatment of drugs, the preincubation period varied depending on the drugs, which were directly added after the resuspension step. Untreated control synaptoneurosomes were pre-incubated for the same period as drug pre-treatment time for comparisons.

### Cell Line and Primary Neuronal Culture

Primary neurons were cultured from cerebral cortices as described by Banker & Goslin, 1998 [84]. Embryos were obtained from females on the 18th day of the gestation period (E18) and cerebral cortices were dissected out in ice-cold Hank’s balanced salt solution under a dissection microscope. Cells were dissociated with 0.25% trypsin solution at 37°C for 10 minutes followed by mechanical trituration in minimal essential medium (MEM, Thermo fisher) with 10% fetal bovine serum (Sigma). Dissociated cells were plated on tissue culture dishes or coverslips coated with poly-L-lysine (0.2 mg/ml in borate buffer, pH 8.5). Neurons were attached to the substrate in MEM with 10% FBS for 3h, followed by defined Neurobasal Medium (Invitrogen) with GlutaMAX™ supplement (Gibco™) and B-27 supplements (Invitrogen) for 15 days at 37°C in a 5% CO2 environment. For immunocytochemistry, cells plated on coverslips kept in a 10cm diameter tissue culture dish with a plating density of 105 cells/dish. The coverslips are then inverted onto an astroglial bed in neurobasal media after substrate attachment in MEM. For biochemical and immunoblotting experiments, cells were grown on 6 well dishes or 35mm diameter dishes, for live imaging cells were plated on 35mm diameter glass-bottom dishes, for immunolabeling cells were plated on 15mm diameter glass coverslips with 0.15mm thickness. The plating density used for various assays are as follows: ATP:ADP Ratio (Biochemical)- 4×10^4^ /Cm^2^ (High-density); ATP:ADP (Imaging)-2×10^4^ /Cm^2^ (Low-density); Western blotting of eEF2-4×10^4^ /Cm^2^ (High-density); FUNCAT, immunolabeling and Ca^2+^ Assay (Imaging)- 2×10^4^ /Cm^2^ (Low-density); KD of α-SNAP and Western Blotting-4×10^4^ /Cm^2^ (High-density).

### Electrophysiology

To test the functionality of the rat cortical neurons whole-cell patch-clamp recordings were performed as previously described [85,86]. Rat cortical neurons were cultured on glass coverslips and transferred onto the recording chamber after 14-15 days *in vitro*. An extracellular solution composed of the following (in mM) NaCl 152, KCl 2.8, HEPES 10, CaCl_2_ 2, glucose 10, pH 7.3 – 7.4 (300 – 320 mOsm) was constantly perfused using a peristaltic pump (Watson – Marlow, Wilmington, Massachusetts, USA). Patch pipettes (3 - 4 MΩ resistance, ≈ 2 µm tip diameter) were pulled from thick-walled Borosilicate glass using a P1000 horizontal micropipette puller (Sutter Instruments, Novato, California, USA. For current-clamp recordings pipettes were filled with an internal solution consisting of the following (in mM) : K-gluconate 155, MgCl_2_ 2, Na-HEPES 10, Na-PiCreatine 10, Mg_2_-ATP 2 and Na_3_-GTP 0.3, pH 7.3 (280 - 290 mOsm) and Cesium gluconate 110, CsCl 20, Na-HEPES 10, NaCl 4, QX 314 5, EGTA 0.2, Na-PiCreatine 10, Mg_2_-ATP 2 and Na_3_-GTP 0.3, pH 7.3 (280 - 290 mOsm) was used for voltage-clamp recordings. Signals were filtered at 3 kHz and 10 kHz for voltage and current clamp, respectively, using a Multiclamp 700B amplifier and digitized at 10 kHz with Axon Digidata 1550 (Molecular Devices, Union City, California, USA). Holding current was less than 100 pA for all the recordings and the series resistance *R*_s_ was less than 30 MΩ. Experiments, where Rs drifted by more than 25%, were discarded. In action potential (AP) recordings, neurons were held at −60 mV and a series of depolarizing current pulses (−40 pA to +540 pA, 500 msec) were injected. AP parameters were analyzed from the first action potential using Clampfit 10.5 (pClamp, Molecular Devices, Union City, California, USA, RRID: SCR_011323). For recording miniature excitatory postsynaptic currents (mEPSC) in the voltage-clamp mode neurons were held at −70mV in the presence of 0.5 µM TTX (Hello Bio, Bristol, UK) and recorded for 5 min of which the last 1 min was used to analyze the mEPSC parameters. For spontaneous burst recordings in both voltage and current clamp, the neurons were clamped at −70 mV and recorded for 5 min. Burst number was defined as the average number of bursts in 5 min and burst duration was calculated as the time interval between the start of membrane depolarization to the end of depolarization. All values are expressed as mean ± SEM. Each data set was tested for normality using the D’Agostino & Pearson test. Column statistics was performed for calculating the mean, standard deviation, and SEM. All the statistical tests were performed using GraphPad (GraphPad Software Inc., La Jolla, California, USA, RRID: SCR_002798).

### Metabolic Labeling

For metabolic labeling of proteins, neurons were incubated for 1 hour in methionine-free Dulbecco’s Modified Essential Medium (DMEM, Thermo Fisher). This was followed by the addition of L-azidohomoalanine (AHA; 1μM) for the next 55 minutes in the same medium. At this point, a group of cells were fixed with 4% Paraformaldehyde (PFA) for 20 minutes and would be considered as unstimulated basal for further analysis. This was followed by either fixing the cells with 4% PFA or the cells were stimulated with mGluR and NMDAR specific agonists DHPG (50uM) and NMDA (20 uM) respectively for 2 minutes, 5 minutes or post-stimulation 5 minutes recovery (5 min stim + 5 min recovery) and subsequently fixed with 4% PFA. Cells were then permeabilized in TBS50 [containing (in mM):50 Tris-Base, 150 NaCl] + 0.3% Triton X 100 solution and blocked using TBS50 + 0.1% Triton X 100 + 2% BSA + 4% FBS containing blocking solution. Newly synthesized proteins were then labeled with Alexa-Fluor-555–alkyne [Alexa Fluor 488 5-carboxamido-(propargyl), bis (triethylammonium salt)], by allowing the fluorophore alkyne to react with AHA azide group through click chemistry. All reagents were from Thermo Fisher and stoichiometry of reagents were calculated according to the suggested manual by the manufacturer (CLICK-iT cell reaction buffer kit, cat.no. C10269). Signal intensity was then measured using confocal microscopy to calculate the amount of newly synthesized proteins. For the detection of neurons, MAP2B immunolabeling was used.

### Immunocytochemistry

Rat primary cortical neurons were stimulated with either 50μM DHPG or 20μM NMDA for 5mins. Cells were fixed with 4% PFA and processed for imaging as described before. In brief, cells were permeabilized using TBS50 + 0.3% T solution and were treated with Tris-Glycine solution (containing in Moles: 0.5 Tris-base and 0.2 Glycine) before blocking with blocking buffer [TBS50 + 0.1% T) + 2% BSA + 4% FBS]. Primary antibodies were incubated in blocking buffer overnight at 4°C under gentle shaking conditions followed by washes. Alexa Fluor 488 coupled anti-mouse and Alexa Fluor 555 coupled anti-rabbit secondary antibodies were incubated for 2h at room temperature. Finally, coverslips were mounted for imaging using Mowiol® 4-88 mounting media. Images were acquired on a Leica TCS SP5 confocal microscope (Leica Biosystems) with HCX PL APO 63X, NA 1.4, oil immersion objective. Imaging conditions were kept constant across groups. Images were acquired on Olympus FLUOVIEW 3000 confocal laser scanning microscope (Olympus Corporation) with HCX PL APO 60x, NA 1.4, oil immersion objective. For the quantification of fluorescence intensities, images were acquired while keeping the open pinhole configuration to collect lights from planes below and above the focal plane as well. The objective was moved in the Z direction with a step size of 0.5μm for 12 such steps to ensure the light was collected from the entire cell across its thickness. For quantification of colocalization, images were acquired using 2.5x optical zooming to satisfy Nyquist’s sampling theorem of information theory for maximum resolution in the XY direction. The pinhole was kept at 1 Airy unit to ensure optimum resolution and confocal stacks were acquired with a step size of 0.3μm calculated based on the fluorophore wavelength. Imaging conditions were kept constant across different data sets, across experiments.

### Live Cell Fluorescence Microscopy

#### Imaging Dendritic ATP/ADP Ratio

Cells were imaged on Zeiss LSM 780 confocal laser scanning microscope (Carl Zeiss, Oberkochen, Germany) with 63X Zeiss plan-apochromatic oil immersion objective, NA 1.4 with argon lasers of specified wavelengths. Cells were grown in Neurobasal media containing B27 supplements and Glutamax and were transfected with PercevalHR and pH-Red construct for simultaneous monitoring of intracellular ATP/ADP ratio and pH. pH monitoring was done as suggested in previous studies (56) to correct for the bias created in Perceval HR fluorescence solely due to the change in intracellular pH. Perceval HR was excited with 488/20 nm and 405/20 nm band-pass filters and emissions were collected through a 520/15nm band-pass filter. Excitation and emission beams were separated through a 490 nm short pass dichroic mirror. pH-Red was excited using 561/20nm and 455/10 nm band-pass filters and emissions were collected through a 630/50 nm band-pass filter. A 560 nm short-pass dichroic was used to separate excitation and emission beams.

Neurons were imaged at room temperature in 37°C pre-warmed Neurobasal media without phenol red (Thermo Fisher, Waltham, MA, USA) containing 15mM HEPES. For, approximate pH bias removal, the linear relationship established between Perceval HR and pH-Red fluorescence for the high glucose ‘ATP Loaded’ state by Tantama et. Al., 2014 [40] was used. The relationship was used to predict the pH bias of the Perceval HR signal from the continuously monitored pH-Red signal and was normalized to the observed Perceval HR fluorescence for the entire period of the experiment. Images were captured at a 16-bit image format with 512×512 pixel distribution with a frame rate of 1 per 15 seconds.

#### Ca2+ Imaging

Growth media from cells grown on glass-bottom Petri dishes were first removed and were washed with imaging media [containing in mM: 120 NaCl, 3 KCl, 2 CaCl2, 1MgCl2, 3 NaHCO3, 1.25 NaH2PO4, 15 HEPES, 30 glucose (pH 7.4). They were incubated with 2 ml freshly prepared dye solution (2 µM Fluo-8 AM and 0.002% Pluronic F-127 in imaging media) at 37°C for 10 mins followed by a 5 minutes incubation procedure at the same temperature with the imaging media. They were then imaged on Olympus FV3000 confocal laser scanning inverted microscope with 20X air objective lens NA 0.75, illuminated with 488nm solid-state lasers. Images were acquired at a rate of single frame per 3.22s intervals at room temperature. Cells were imaged for 322 sec (100 frames) for recording spontaneous activity, followed by 644 sec (200 frames) with stimulant and 161 sec (50 frames) with KCl (to check neuronal activity) or ionomycin (Fmax). After background, fluorescence were calibrated to [Ca2+]i as [Ca2+]i = 389 (F-Fmin)/(Fmax-F)nM. Fmin was recorded by chelating Ca2+ with 10mM BAPTA (Sigma) in independent experiments and Fmax was recorded by 10mM ionomycin in the presence of 10mM CaCl_2_ in each experiment.

#### Luciferase Assay

To measure synaptic ATP levels, Cortical synaptoneurosomes were first pre-warmed and then stimulated with Gp I mGluR and NMDAR specific agonists at 37°C under constant shaking. Fractions of the stimulated solutions were collected at the time points specified before. For treatment with various drugs, the pre-warming period varied depending on the pre-incubation time of the drug before the stimulants were added. Collected fractions were added to an equal volume of boiling water to extract the ATP as described previously (Yang et al., 2002). The lysates were then used for quantification of soluble ATP molecules using luciferase-based commercial ATP quantification kit (ATP Determination kit, Thermo Fisher, Cat.no. A22066) with the help of a standard curve. For measuring ATP/ADP ratio from cortical neurons, cells were stimulated for specified time points and lysed with lysis buffer [containing in mM: 50 Tris-Cl (pH-7.4), 150 NaCl, 5 MgCl2, 1% Triton-X-100, supplemented with EDTA free protease inhibitor complex (Sigma, cat. no. S8830) and phosphatase inhibitor cocktail (Roche, ref.no. 04906837001)]. Lysates were then centrifuged at 20,000g for 20 minutes and supernatants were used for measuring ATP using luciferase-based protocol. This was followed by a step converting ADP to ATP which was then used to measure the ATP and ADP level together constituting the bulk of the adenine nucleotides. This was done using a commercially available luminescence-based kit (ADP/ATP ratio kit, Abcam, cat.no. ab65313) following the user manual suggested by the manufacturer.

#### Image Analysis

Image analysis was done using Fiji (ImageJ based image processing package) and IMARIS 9.0 (Bitplane, Oxford instrument company, Belfast, UK) software. For fluorescence intensity quantification, confocal stacks were collapsed using the sum slice projection method and mean intensity was quantified from the cells. MAP2B positive cells were used for the identification of neurons and quantified mean intensity values were used for normalization. For quantifying colocalization, Pearson’s correlation coefficient (PCC) was measured for the cell body and proximal dendrite region (<50μm). For the coefficient analysis, the measurement was done after manual thresholding with a threshold value of 155 for a 12bit image format. For time frame analysis, mean fluorescence intensity was quantified for the imaged dendritic region from each time frame. Time frame analysis was done using the time series analyzer V3 plug-in from Fiji. Representative images for ratiometric quantifications were generated using the Ratio Plus plug from Fiji where the image was generated by calculating pixel by pixel intensity ratio between two channels.

### Statistical Analysis

Statistical comparisons were done using GraphPad Prism (Prism 7.01, GraphPad Software Inc, La Jolla, CA, USA). For group comparisons, the data distributions were first tested for normality using the Kolmogorov-Smirnov Goodness-of-Fit Test or Shapiro-Wilk normality test. Depending on the distribution, either parametric or non-parametric tests were used to quantify statistical significance. For groups with <4 data points, the inherent distribution was considered to follow Gaussian distribution unless mentioned. For comparing two groups, Student’s t-test (two-tailed, paired/unpaired) was done to calculate statistical significance. Multiple group comparisons were made using one-way ANOVA followed by Bonferroni’s multiple comparison test for parametric distribution and Kruskal-Wallis’s test followed by Dunn’s multiple comparison test. Two-way ANOVA followed by Holm Sidak’s multiple comparison test was performed for comparing multiple groups at various time points. For calculating the variance of the basal ATP/(ATP+ADP) ratio, values obtained from different plates from multiple cell culture experiments were considered. For calculating the variance of basal synaptic ATP level at 0 min, synaptoneurosomal ATP content quantified from multiple animals were considered. For calculating the variance of ATP/ADP ratio at 0 min in live imaging experiments, values obtained from neurons from multiple culture experiments were considered. For calculating the basal fluorescence intensity variance in the immunolabelling and FUNCAT experiments, intensity values obtained from different cells obtained from multiple culture experiments were considered. For calculating the phospho/total ratio of a protein, the phospho form, and the total protein was probed in individual blots and normalized to their corresponding tubulin levels for every time point. Further, a ratio of the two normalized values (i.e. normalized phospho/normalized total) was considered as the phospho/total ratio. The ratio for all the time points are expressed as a fraction of the initial 0 min value. Data are presented as mean ± SEM in the case of bar graphs. Data presented in box plots show the distribution of all data points for the group including minimum and maximum values. The box extends from 25^th^ to 75^th^ percentile with the middlemost line representing the median of the dataset. Whiskers range from minimum to maximum data point. *p< 0.05 was considered statistically significant.

## Acknowledgment

We are thankful to the Central Imaging and Flow-cytometry Facility (CIFF), NCBS for allowing us to use the LSM 780 and FV3000 confocal microscopes, NCBS animal house facility for generously maintaining the rat colony needed for experimentation. We extend our gratitude to Professor M.K. Mathew and Dr. Tina Mukherjee for their helpful comments and discussions. We thank Chaitra Ananda, Amirthavarshini Devarajan, and Sonu P. Kurian for their gracious efforts in experimental support and data analysis. The work was funded by the NeuroStem grant (BT/IN/Denmark/07/RSM/2015-2016), Center for Brain Development and Repair (CBDR) (BT/MB-CNDS/2013) and the Department of Biotechnology, Govt. of India. We thank Dr. Garry Yellen, Harvard University for his useful suggestions on PercevalHR experiments.

## Author contribution

SGD designed and performed the experiments, analyzed data and co-wrote the manuscript, SDS performed Ephys experiments and analyzed data, SCB performed Calcium imaging experiments and analyzed data, SC provided resources, AB provided resources, analyzed data and co-wrote the manuscript, RM conceptualized the question, designed experiments, provided resources, and co-wrote the manuscript.

## References

1. Mink JW, Blumenschine RJ, Adams DB. Ratio of central nervous system to body metabolism in vertebrates: its constancy and functional basis. Am J Physiol Integr Comp Physiol. 1981;241(3):R203–12.

2. Fox PT, Raichle ME. Focal physiological uncoupling of cerebral blood flow and oxidative metabolism during somatosensory stimulation in human subjects. Proc Natl Acad Sci U S A. 1986;83(4):1140–4.

3. Fox PT, Mintun MA, Reiman EM, Raichle ME. Enhanced detection of focal brain responses using intersubject averaging and change-distribution analysis of subtracted PET images. J Cereb Blood Flow Metab. 1988;8(5):642–53.

4. Attwell D, Laughlin SB. An energy budget for signaling in the grey matter of the brain. J Cereb Blood Flow Metab. 2001;21(10):1133–45.

5. Harris JJ, Jolivet R, Attwell D. Synaptic energy use and supply. Neuron. 2012;75(5):762–77.

6. Gafaniz R, Sanches JM. ATP consumption and neural electrical activity: a physiological model for brain imaging. 2010 Annu Int Conf IEEE Eng Med Biol Soc EMBC’10. 2010;5480–3.

7. Zhu F, Wang R, Pan X, Zhu Z. Energy expenditure computation of a single bursting neuron. Cogn Neurodyn. 2019;13(1):75–87.

8. Misgeld T, Schwarz TL. Mitostasis in neurons: maintaining mitochondria in an extended cellular architecture. Neuron. 2017;96(3):651–66.

9. Rangaraju V, Calloway N, Ryan TA. Activity-driven local ATP synthesis is required for synaptic function. Cell. 2014;156(4):825–35.

10. Pathak D, Shields LY, Mendelsohn BA, Haddad D, Lin W, Gerencser AA, Kim H, Brand MD, Edwards RH, Nakamura K. The role of mitochondrially derived ATP in synaptic vesicle recycling. J Biol Chem. 2015;290(37):22325–36.

11. Hsu WL, Chung HW, Wu CY, Wu HI, Lee YT, Chen EC, Fang W, Chang YC. Glutamate stimulates local protein synthesis in the axons of rat cortical neurons by activating α-amino-3-hydroxy-5-methyl-4-isoxazolepropionic acid (AMPA) receptors and metabotropic glutamate receptors. J Biol Chem. 2015;290(34):20748–60.

12. Kos A, Wanke KA, Gioio A, Martens GJ, Kaplan BB, Aschrafi A. Monitoring mrna translation in neuronal processes using fluorescent non-canonical amino acid tagging. J Histochem Cytochem. 2016;64(5):323–33.

13. Rangaraju V, Lauterbach M, Schuman EM. Spatially stable mitochondrial compartments fuel local translation during plasticity. Cell. 2019;176(1–2):73–84.e15.

14. Rangaraju V, tom Dieck S, Schuman EM. Local translation in neuronal compartments: how local is local? EMBO Rep. 2017;18(5):693–711.

15. Mery F, Kawecki TJ. A cost of long-term memory in drosophila. Science (80-). 2005;308(5725):1148.

16. Plaçais PY, Preat T. To favor survival under food shortage, the brain disables costly memory. Science (80-). 2013;339(6118):440–2.

17. Abraham WC, Williams JM. LTP maintenance and its protein synthesis-dependence. Neurobiol Learn Mem. 2008;89(3):260–8.

18. Schwanhüusser B, Busse D, Li N, Dittmar G, Schuchhardt J, Wolf J, Chen W, Selbach M. Global quantification of mammalian gene expression control. Nature. 2011;473(7347):337–42.

19. Huber KM, Kayser MS, Bear MF. Role for rapid dendritic protein synthesis in hippocampal depression. Science (80-). 2000;288(May):1254–6.

20. Luchelli L, Thomas MG, Boccaccio GL. Synaptic control of mRNA translation by reversible assembly of Xrn1 bodies. J Cell Sci. 2015;128(8):1542–54.

21. Scheetz AJ, Nairn AC, Constantine-paton M. NMDA receptor-mediated control of protein synthesis at developing synapses. Nat Neurosci. 2000;211–6.

22. Hunt DL, Castillo PE. Synaptic plasticity of NMDA receptors: mechanisms and functional implications. Curr Opin Neurobiol. 2012;22(3):496–508.

23. Malenka L. NMDA receptor-dependent long-term potentiation and long-term depression (LTP/LTD). Cold Spring Harb Perspect Biol. 2012;1–16.

24. Bear MF, Huber KM, Warren ST. The mGluR theory of fragile x mental retardation. Trends Neurosci. 2004;27(7):370–7.

25. Marinangeli C, Didier S, Ahmed T, Caillerez R, Domise M, Laloux C, Bégard S, Carrier S, Colin M, Marchetti P, et al. AMP-activated protein kinase is essential for the maintenance of energy levels during synaptic activation. IScience. 2018;9:1–13.

26. Manlio Díaz-García C, Mongeon R, Lahmann C, Koveal D, Zucker H, Correspondence GY, Yellen G. Neuronal stimulation triggers neuronal glycolysis and not lactate uptake cell metabolism article neuronal stimulation triggers neuronal glycolysis and not lactate uptake. Cell Metab. 2017;26:361–374.e4.

27. Bélanger M, Allaman I, Magistretti PJ. Brain energy metabolism: focus on astrocyte-neuron metabolic cooperation. Cell Metab. 2011;14(6):724–38.

28. Burkewitz K, Zhang Y, Mair WB. AMPK at the nexus of energetics and aging. Cell Metab. 2014;20(1):10–25.

29. Potter WB, O’Riordan KJ, Barnett D, Osting SMK, Wagoner M, Burger C, Roopra A. Metabolic regulation of neuronal plasticity by the energy sensor AMPK. PLoS One. 2010;5(2).

30. Kong D, Dagon Y, Campbell JN, Guo Y, Yang Z, Yi X, Aryal P, Wellenstein K, Kahn BB, Sabatini BL, et al. A postsynaptic AMPK?p21-activated kinase pathway drives fasting-induced synaptic plasticity in AgRP neurons. Neuron. 2016;91(1):25–33.

31. Sutton MA, Wall NR, Aakalu GN, Schuman EM. Regulation of dendritic protein synthesis by miniature synaptic events. Science (80-). 2004;304(5679):1979–83.

32. Fishbein I, Segal M. Miniature synaptic currents become neurotoxic to chronically silenced neurons. Cereb Cortex. 2007;17(6):1292–306.

33. Zhu G, Du L, Jin L, Offenhäusser A. Effects of morphology constraint on electrophysiological properties of cortical neurons. Sci Rep. 2016;6(February):1–10.

34. Dichter MA. Rat cortical neurons in cell culture: culture methods, cell morphology, electrophysiology, and synapse formation. Brain Res. 1978;149(2):279–93.

35. Neumann JR, Dash-Wagh S, Jack A, Räk A, Jüngling K, Hamad MIK, Pape HC, Kreutz MR, Puskarjov M, Wahle P. The primate-specific peptide Y-p30 regulates morphological maturation of neocortical dendritic spines. PLoS One. 2019;14(2):1–23.

36. Dichter MA, Lisak J, Biales B. Action potential mechanism of mammalian cortical neurons in cell culture. Brain Res. 1983;289(1–2):99–107.

37. Charlesworth P, Cotterill E, Morton A, Grant SGN, Eglen SJ. Quantitative differences in developmental profiles of spontaneous activity in cortical and hippocampal cultures. Neural Dev. 2015;10(1):1–10.

38. Kute PM, Ramakrishna S, Neelagandan N, Chattarji S, Muddashetty RS. NMDAR mediated translation at the synapse is regulated by MOV10 and FMRP. Mol Brain. 2019;12(1):1–14.

39. Cottrell JR, Dubé GR, Egles C, Liu G. Distribution, density, and clustering of functional glutamate receptors before and after synaptogenesis in hippocampal neurons. J Neurophysiol. 2000;84(3):1573–87.

40. Tantama M, Martínez-François JR, Mongeon R, Yellen G. Imaging energy status in live cells with a fluorescent biosensor of the intracellular ATP-to-ADP ratio. Nat Commun. 2013;4(May).

41. Pedersen SF, Jørgensen NK, Damgaard I, Schousboe A, Hoffmann EK. Mechanisms of pH(i) regulation studied in individual neurons cultured from mouse cerebral cortex. J Neurosci Res. 1998;51(4):431–41.

42. Biffi E, Regalia G, Menegon A, Ferrigno G, Pedrocchi A. The influence of neuronal density and maturation on network activity of hippocampal cell cultures: a methodological study. PLoS One. 2013;8(12).

43. Ivenshitz M, Segal M. Neuronal density determines network connectivity and spontaneous activity in cultured hippocampus. J Neurophysiol. 2010;104(2):1052–60.

44. Mangan PS, Kapur J. Factors underlying bursting behavior in a network of cultured hippocampal neurons exposed to zero magnesium. J Neurophysiol. 2004;91(2):946–57.

45. Vergara RC, Jaramillo-Riveri S, Luarte A, Moënne-Loccoz C, Fuentes R, Couve A, Maldonado PE. The energy homeostasis principle: neuronal energy regulation drives local network dynamics generating behavior. Front Comput Neurosci. 2019;13(July):1–18.

46. Dieterich DC, Hodas JJL, Gouzer G, Shadrin IY, Ngo JT, Triller A, Tirrell DA, Schuman EM. In situ visualization and dynamics of newly synthesized proteins in rat hippocampal neurons. Nat Neurosci. 2010;13(7):897–905.

47. Ferrara NC, Jarome TJ, Cullen PK, Orsi SA, Kwapis JL, Trask S, Pullins SE, Helmstetter FJ. GluR2 endocytosis-dependent protein degradation in the amygdala mediates memory updating. Sci Rep. 2019;9(1):1–10.

48. Sheehan P, Zhu M, Beskow A, Vollmer C, Waites CL. Activity-dependent degradation of synaptic vesicle proteins requires Rab35 and the ESCRT pathway. J Neurosci. 2016;36(33):8668–86.

49. Jarome TJ, Werner CT, Kwapis JL, Helmstetter FJ. Activity dependent protein degradation is critical for the formation and stability of fear memory in the amygdala. PLoS One. 2011;6(9).

50. Richter JD, Coller J. Pausing on polyribosomes: make way for elongation in translational control. Cell. 2015;163(2):292–300.

51. Maus M, Torrens Y, Gauchy C, Bretin S, Nairn AC, Glowinski J, Premont J. 2- deoxyglucose and NMDA inhibit protein synthesis in neurons and regulate phosphorylation of elongation factor-2 by distinct mechanisms. J Neurochem. 2006;96(3):815–24.

52. Ryazanov Alexey G., Shestakova Elena A. NPG. Phosphorylation of elongation factor 2 by EF2-kinase affects the rate of translation. Nature. 1988;334(14):170–3.

53. Nairn AC, Bhagat B, Palfrey HC. Identification of calmodulin-dependent protein kinase iii and its major m(r) 100,000 substrate in mammalian tissues. Proc Natl Acad Sci U S A. 1985;82(23):7939–43.

54. Richter JD. RNA and the synapse. Rna. 2015;21(4):716–7.

55. Ryazanov AG, Davydova EK. Mechanism of elongation factor 2 (EF-2) inactivation upon phosphorylation: phosphorylated EF-2 is unable to catalyze translocation. FEBS Lett. 1989;251(1–2):187–90.

56. Takei N, Kawamura M, Ishizuka Y, Kakiya N, Inamura N, Namba H, Nawa H. Brain-derived neurotrophic factor enhances the basal rate of protein synthesis by increasing active eukaryotic elongation factor 2 levels and promoting translation elongation in cortical neurons. J Biol Chem. 2009;284(39):26340–8.

57. Browne GJ, Finn SG, Proud CG. Stimulation of the AMP-activated protein kinase leads to activation of eukaryotic elongation factor 2 kinase and to its phosphorylation at a novel site, serine 398. J Biol Chem. 2004;279(13):12220–31.

58. Hawley SA, Davison M, Woods A, Davies SP, Beri RK, Carling D, Hardie DG. Characterization of the AMP-activated protein kinase kinase from rat liver and identification of threonine 172 as the major site at which it phosphorylates AMP-activated protein kinase. J Biol Chem. 1996;271(44):27879–87.

59. Joseph BK, Liu HY, Francisco J, Pandya D, Donigan M, Gallo-Ebert C, Giordano C, Bata A, Nickels JT. Inhibition of AMP kinase by the protein phosphatase 2a heterotrimer, pp2appp2r2d. J Biol Chem. 2015;290(17):10588–98.

60. Voss M, Paterson J, Kelsall IR, Martín-Granados C, Hastie CJ, Peggie MW, Cohen PTW. Ppm1E is an in cellulo AMP-activated protein kinase phosphatase. Cell Signal. 2011;23(1):114–24.

61. Wang L, Brautigan DL. α-snap inhibits AMPK signaling to reduce mitochondrial biogenesis and dephosphorylates Thr172 in AMPKα in vitro. Nat Commun. 2013;4:1559.

62. Woods A, Dickerson K, Heath R, Hong SP, Momcilovic M, Johnstone SR, Carlson M, Carling D. Ca2+/calmodulin-dependent protein kinase kinase-β acts upstream of AMP-activated protein kinase in mammalian cells. Cell Metab. 2005;2(1):21–33.

63. Maus M, Marin P, Israël M, Glowinski J, Prémont J. Pyruvate and lactate protect striatal neurons against n-methyl-d-aspartate-induced neurotoxicity. Eur J Neurosci. 1999;11(9):3215–24.

64. Kim J, Yang G, Kim Y, Kim J, Ha J. AMPK activators: mechanisms of action and physiological activities. Exp Mol Med. 2016;48(4):1–12.

65. Yoon Byung C., Zivraj Krishna H, Holt CE. Local translation and mRNA trafficking in axon pathfinding. Results Probl Cell Differ 2009;48:269–88.

66. Spillane M, Ketschek A, Merianda TT, Twiss JL, Gallo G. Mitochondria coordinate sites of axon branching through localized intra-axonal protein synthesis. Cell Rep. 2013;5(6):1564–75.

67. Hovy Ho-Wai Wong A, Qiaojin Lin J, Ströhl F, Harris WA, Kaminski CF, Holt CE, Ho-Wai Wong H, udio Gouveia Roque C, Cioni J-M, Cagnetta R, et al. RNA docking and local translation regulate site-specific axon remodeling in vivo article RNA docking and local translation regulate site-specific axon remodeling in vivo. Neuron. 2017;95:852–68.

68. Cajigas J, Will T, Schuman EM. Protein homeostasis and synaptic plasticity. 2010;29(16):2746–52.

69. Rosenberg T, Gal-ben-ari S, Dieterich DC, Kreutz MR, Ziv NE. The roles of protein expression in synaptic plasticity and memory consolidation. 2014;7(November):1–14.

70. Nödling AR, Spear LA, Williams TL, Luk LYP, Tsai YH. Using genetically incorporated unnatural amino acids to control protein functions in mammalian cells. Essays Biochem. 2019;63(2):237–66.

71. Alvarez-Castelao B, Schuman EM. The regulation of synaptic protein turnover. J Biol Chem. 2015;290(48):28623–30.

72. Weiler IJ, Spangler CC, Klintsova AY, Grossman AW, Kim SH, Bertaina-Anglade V, Khaliq H, de Vries FE, Lambers F a E, Hatia F, et al. Fragile x mental retardation protein is necessary for neurotransmitter-activated protein translation at synapses. Proc Natl Acad Sci USA. 2004;101(50):17504–9.

73. Bellone C, Lüscher C, Mameli M. Mechanisms of synaptic depression triggered by metabotropic glutamate receptors. Cell Mol Life Sci. 2008;65(18):2913–23.

74. Schulte G. The class frizzled receptors. Pharmacol Rev. 2010;62(4):632–67.

75. Jörntell H, Hansel C. Synaptic memories upside down: bidirectional plasticity at cerebellar parallel fiber-purkinje cell synapses. Neuron. 2006;52(2):227–38.

76. Yuan Y, Huo H, Fang T. Effects of metabolic energy on synaptic transmission and dendritic integration in pyramidal neurons. Front Comput Neurosci. 2018;12(September):1–13.

77. Peth A, Nathan JA, Goldberg AL. The ATP costs and time required to degrade ubiquitinated proteins by the 26s proteasome. J Biol Chem. 2013;288(40):29215–22.

78. Ashrafi G, Wu Z, Farrell RJ, Ryan TA. GLUT4 mobilization supports energetic demands of active synapses. Neuron. 2017;93(3):606–615.e3.

79. Johanns M, Pyr dit Ruys S, Houddane A, Vertommen D, Herinckx G, Hue L, Proud CG, Rider MH. Direct and indirect activation of eukaryotic elongation factor 2 kinase by AMP-activated protein kinase. Cell Signal. 2017;36(April):212–21.

80. Narayanan U, Nalavadi V, Nakamoto M, Pallas DC, Ceman S, Bassell GJ, Warren ST. FMRP phosphorylation reveals an immediate-early signaling pathway triggered by group i mGluR and mediated by PP2a. J Neurosci. 2007;27(52):14349–57.

81. Hollingsworth EB, McNeal ET, Burton JL, Williams RJ, Daly JW, Creveling CR. Biochemical characterization of a filtered synaptoneurosome preparation from guinea pig cerebral cortex: cyclic adenosine 3’:5’-monophosphate-generating systems, receptors, and enzymes. J Neurosci. 1985;5(8):2240–53.

82. Nalavadi VC, Muddashetty RS, Gross C, Bassell GJ. Dephosphorylation-induced ubiquitination and degradation of FMRP in dendrites: a role in immediate early mGluR-stimulated translation. J Neurosci. 2012;32(8):2582–7.

83. Yang NC, Ho WM, Chen YH, Hu ML. A convenient one-step extraction of cellular ATP using boiling water for the luciferin-luciferase assay of atp. Anal Biochem. 2002;306(2):323–7.

84. Banker G, Goslin K. Culturing nerve cells. In: Cellular and molecular neuroscience series. 1998. p. 666.

85. Bilican B, Livesey MR, Haghi G, Qiu J, Burr K, Siller R, Hardingham GE, Wyllie DJA, Chandran S. Physiological normoxia and absence of EGF is required for the long-term propagation of anterior neural precursors from human pluripotent cells. PLoS One. 2014;9(1).

86. Livesey MR, Bilican B, Qiu J, Rzechorzek NM, Haghi G, Burr K, Hardingham GE, Chandran S, Wyllie DJA. Maturation of AMPAR composition and the gabaar reversal potential in hpsc-derived cortical neurons. J Neurosci. 2014;34(11):4070–5.

