## Supplementary Figure Legends for "Distinct Regulation of Bioenergetics and Translation by Group I mGluR and NMDAR"

**Figure EV 1:**

**Quantification of activity-dependent changes in the energy level** (Related to Figure **(A)** Bar graph representing the normalized average value of the neuronal ATP/ATP+ADP ratio in basal condition, on DHPG (50μM) and on NMDA (20μM) treatment for 5 minutes. Data presented as mean ± SEM. *p<0.05. Values in all other groups were normalized to the basal group for the corresponding experiment. n=3 independent plating. One-way ANOVA followed by Bonferroni’s multiple comparison test.

**(B)** Bar graph representing the normalized average value of the neuronal ATP/ATP+ADP ratio in basal condition, on Glutamate (25μM) treatment for 5 minutes and on glutamate treatment with dynaosre (100μM). Dynasore was pre-incubated for 30 minutes. Data presented as mean ± SEM. Values in all other groups were normalized to the basal group for the corresponding experiment. *p<0.05, n=4 independent plating. One-way ANOVA followed by Bonferroni’s multiple comparison test.

**(C)** Bar graph representing the normalized average value of the neuronal ATP/ATP+ADP ratio in basal condition, on Glutamate (25μM) treatment for 5 minutes, on glutamate treatment with ouabain (1mM) and on glutamate treatment with TTX (1μM). Ouabain and TTX were pre-incubated for 3 minutes. Data presented as mean ± SEM. Values in all other groups were normalized to the basal group for the corresponding experiment. *p<0.05, n=5 independent plating. One-way ANOVA followed by Bonferroni’s multiple comparison test.

**(D)** Bar graph representing the effect of individual drugs alone over the neuronal ATP/ATP+ADP ratio in neurobasal media. All the drugs were incubated for 30 minutes before glutamate addition (figure 1B, 1C and Figure EV 1B). Data presented as mean ± SEM. Values in all other groups were normalized to the basal group for the corresponding experiment. The effects of the drugs were tested using the One-way ANOVA.

**(E)** Bar graph representing the effect of ouabain and TTX alone over the neuronal ATP/ATP+ADP ratio in ACSF. All the drugs were incubated for 3 minutes as per the pre-treatment conditions before glutamate stimulation was presented in Figure EV 1C. Data presented as mean ± SEM. Values in all other groups were normalized to the basal group for the corresponding experiment. n=5 independent platings. The effects of the drugs were tested using the One-way ANOVA.

**(F)** Representative transmission electron micrograph image of the cortical synaptoneurosome preparations used for quantification of activity-dependent changes on various parameters. The preparation shows enrichment of ‘snow-man’ shaped pre and post-synaptic compartments. Scale bar 200nm.

**(G)** Bar graph representing the effect of individual drug treatments over the synaptic ATP level compared to untreated basal conditions. All the drugs were incubated for 10 minutes as per the pre-treatment conditions before DHPG or NMDA stimulation was presented in figure 1H and 1I. Data presented as mean ± SEM. Values in all other groups were normalized to the basal group for the corresponding experiment. n=3 animals. The effects of the drugs were tested using the One-way ANOVA.

**(H)** Representative current-clamp traces showing single action potential fired in response to rheobase current (*top*, 500 ms, +80 pA) and multiple action potentials fired at a higher potential (*bottom*, 500ms, +180 pA) by a 15 days old neuron plated at high-density.

**(I)** Active membrane properties of the neurons: spike amplitude (88.69 ± 3.828 mV), spike half-width (1.862 ± 0.1392 ms), spike threshold (-46.81 ± 2.28 mV).

**(J)** Passive membrane properties of the neurons: capacitance (59.95 ± 4.844 pF), input resistance (184.6 ± 24.54 MΩ) and resting membrane potential (-57.68 ± 1.381 mV).

**(K)** Firing frequency of the high-density neurons calculated as an average number of spikes fired across a range of current injections. n=14 neurons from ≥3 independent platings.

**(L)** Firing frequency of the high-density neurons calculated as an average number of spikes fired across a range of current injections. n=12 neurons from ≥3 independent platings.

**(M)** Bar graph representing the normalized average value of the ATP/ATP+ADP ratio on the basal condition, on glutamate stimulation for 5 minutes, on glutamate stimulation with anisomycin (25μM) from neurons used in the batch of electrophysiology experiments. Data presented as mean ± SEM with scattered data points. Values in all other groups were normalized to the basal group for the corresponding experiment. *p<0.05, **p<0.01, n=7 independent platings. One-way ANOVA followed by Bonferroni’s multiple comparison test.

**Figure EV 2:**

**Sensor Calibration and Basal ATP measurement** (Related to Figure 2)

**(A)** Line Graph depicting a linear relationship established between the PercevalHR fluorescence ratio and the pH-Red fluorescence ratio using NH_4_Cl pre-pulse method as described previously (Pederson et. al, 1998, Tantama et. al, 2013). Briefly, a gradient of 5-15mM NH_4_Cl solution was applied to allow the changes in intracellular pH in a short period to avoid any metabolic stress. Changes in PercevalHR fluorescence were measured along with pH-Red fluorescence and the linear relation was used to remove the pH bias approximately.

**(B)** Representative soma and dendrite showing the temporal profile of PercevalHR fluorescence (ATP/ADP) in unstimulated conditions. Scale bar 10μm.

**(C)** Bar graphs representing the average change in the PercevalHR fluorescence (ΔF/F) on basal conditions, on DHPG treatment for 2 minutes, on NMDA treatment for 2 minutes, on DHPG treatment with anisomycin for 2 minutes and on NMDA treatment with anisomycin for 2 minutes. Data presented as +/- SEM. *p<0.05, **p<0.01, n= 6-9 cells from ≥3 independent platings. One Way ANOVA followed by Bonferroni’s multiple comparison test.

**(D)** Bar graphs representing the average change in the PercevalHR fluorescence (ΔF/F) on basal conditions, on DHPG treatment for 5 minutes, on NMDA treatment for 5 minutes, on DHPG treatment with anisomycin for 5 minutes and on NMDA treatment with anisomycin for 5 minutes. Data presented as mean +/- SEM. *p<0.05, **p<0.01, n= 6-9 cells from ≥3 independent plating. One Way ANOVA followed by Bonferroni’s multiple comparison test.

**(E)** Bar graphs representing the average change in the PercevalHR fluorescence (ΔF/F) on basal conditions, on DHPG treatment for 10 minutes, on NMDA treatment for 10 minutes, on DHPG treatment with anisomycin for 10 minutes and on NMDA treatment with anisomycin for 10 minutes. Data presented as mean +/- SEM. *p<0.05, **p<0.01, n= 6-9 cells from ≥3 independent plating. Kruskal Walli’s test followed by Dunn’s multiple comparison test.

**(F)** Bar graphs representing the comparison of the PercevalHR fluorescence change (ΔF/F) between untreated and anisomycin treated conditions (30 minutes of pre-incubation + 4 minutes of baseline imaging) i.e. t=0 min. Data presented as mean ± SEM. *p<0.05, n≥7 cells from ≥5 independent platings. Unpaired-sample t-test.

**(G)** Bar graphs representing the change in the PercevalHR fluorescence (ΔF/F) at various time-points after anisomycin treatment. Data presented as mean ± SEM. n=7 cells from ≥5 independent platings.

**(H)** Normalized average traces showing the time course of the synaptic ATP levels on unstimulated conditions in cortical synaptoneurosomes. Data presented as mean ± SEM. Values in all other groups were normalized to the 0 min group for the corresponding experiment and all the data points are represented as a fraction of the initial time point. n=5 animals.

**(I)** Representative dendrites showing normalized surface mGluR5 levels on basal and anisomycin treated conditions. The values were normalized to corresponding MAP2B intensities. The associated box plot shows the quantification of the normalized surface mGluR5 level on basal and anisomycin treated conditions. Data points in all groups were normalized to the average of the basal group. n≥28 cells from 3 independent platings.

**(J)** Representative dendrites showing surface NR1 levels on basal and anisomycin treated conditions. The values were normalized to corresponding MAP2B intensities. The associated box plot shows the quantification of the normalized surface NR1 level on basal and anisomycin treated conditions. Data points in all groups were normalized to the average of the basal group. n≥31 cells from 4 independent platings, Unpaired-sample t-test with Welch’s correction.

**(K)** Normalized average traces depicting the time course of the dendritic ATP/ADP ratio on Glutamate stimulation in the presence of glucose and deoxy-glucose. In the pre-stimulation phase, the ATP/ADP ratio was quantified in the presence of glucose or deoxy-glucose at the baseline following which the cells were stimulated with glutamate. Data presented as mean +/- SEM, n= 10 dendrites from 6 cells from 3 independent platings. Values in all other groups were normalized to the 0 min group for the corresponding experiment and all the data points are represented as a fraction of the initial time point.

**(L)** Bar graphs representing a comparison of the PercevalHR fluorescence change (ΔF/F) following 10 minutes of either glucose or deoxy-glucose incubation without stimulation in cultured cortical neurons DIV 15. Data presented as mean ± SEM. ****p<0.0001 n= 10 dendrites from 6 cells from 3 independent plating. Unpaired-sample t-test.

**(M)** Bar graphs representing a comparison of the PercevalHR fluorescence change (ΔF/F) following 1.25 minutes of glutamate stimulation in the presence of either glucose or deoxy-glucose in cultured cortical neurons DIV 15. Data presented as mean ± SEM. n= 10 dendrites from 6 cells from 3 independent platings.

**(N)** Bar graphs representing a comparison of the PercevalHR fluorescence change (ΔF/F) following 5 minutes of glutamate stimulation in the presence of either glucose or deoxy-glucose in cultured cortical neurons DIV 15. Data presented as mean ± SEM. n= 10 dendrites from 6 cells from 3 independent platings.

**Figure EV 3:**

**Translation Kinetics** (Related to Figure 3)

**(A)** Representative images showing newly synthesized proteins visualized through FUNCAT metabolic labeling (pseudo-colored) in cortical neurons at various time points in unstimulated condition. Metabolic labeling in the absence of AHA was used to test the specificity of labeling new proteins. MAP2B immunolabeling (green, inset) was used for identifying neurons and intensity was used for normalization. The average trace quantification and data point distribution for unstimulated cells have been presented in Figure 3C and 3D. Scale bar 10μm.

**(B)** Box plot showing the FUNCAT intensity distribution across multiple neurons shown corresponding to the line graph presented in figure 3D. Data presented as mean +/- SEM. Data points in all groups were normalized to the average of the basal group at 0 min. *p<0.05 **p<0.01 ***p<0.001 ****p<0.001, n= 21-54 neurons per group from 3 independent platings. One Way ANOVA followed by Bonferroni’s multiple comparison tests. B0= at 0 min basal, B2= at 2 min basal, B5= at 5 min basal, B5R= basal at 5 mins of recovery, D2= at 2 min after DHPG treatment, D5= at 5 min after DHPG treatment, D5R= at 5 min of recovery after DHPG treatment, N2= at 2 min after NMDA treatment, N5= at 5 min after NMDA treatment, N5R= at 5 mins of recovery after NMDA treatment.

**(C)** Representative images showing newly synthesized proteins visualized through FUNCAT metabolic labeling (pseudo-colored) in cortical neurons at basal condition, after 2 minutes of NMDA (20μM) treatment, after 20 minutes of NMDA treatment, unstimulated basal in presence of proteasome inhibitor Mg132 and 2 minutes after NMDA stimulation in presence of Mg132. MAP2B immunolabeling was used for identifying neurons and intensity was used for normalization. Scale bar 10μm.

**(D)** Box plot showing the FUNCAT intensity distribution across multiple neurons basal (N0), after 2 minutes of NMDA (20μM) treatment (N2), after 20 minutes of NMDA treatment (N20), unstimulated basal in presence of proteasome inhibitor Mg132 and 2 minutes after NMDA stimulation in presence of Mg132 (N2+Mg132). Data presented as mean +/- SEM. Data points in all groups were normalized to the average of the basal group. *p<0.05 n= 19-40 neurons per group from 3 independent plating. Kruskal Walli’s test followed by Dunn’s multiple comparison test.

**(E)** Representative immunoblots describing changes in phospho-eEF2 and total-eEF2 levels at various time points after DHPG (50μM) and NMDA (20μM) treatment to cultured cortical neurons. Note in each case phospho and total eEF2 levels were normalized individually to tubulin before calculating the ratio.

**(F)** Line graph showing the average value for neuronal phospho/total ratio of eEF2 at various time points after DHPG and NMDA treatment and 5 minutes and 10 minutes after recovery. Data presented as mean +/- SEM. Values in all other groups were normalized to the 0 min group for the corresponding experiment and all the data points are represented as a fraction of the initial time point. *p<0.05, **p<0.01, ***p<0.001, n= 3 animals per group. Two Way ANOVA followed by Bonferroni’s multiple comparison test.

**Figure EV 4:**

**Baseline eEF2 phosphorylation and Characterization of NMDAR dependent Ca^2+^ entry** (Related to Figure 4 and Figure 5)

**(A)** Line graph showing the average value of phospho/total eEF2, phospho/total AMPK and α-SNAP at various time points on the basal condition in cortical synaptoneurosomes. Data presented as mean +/- SEM. Values in all other groups were normalized to the 0 min group for the corresponding experiment and all the data points are represented as a fraction of the initial time point. N=3 animals per group.

(B) Line graph showing the average value for phospho/total eEF2, phospho/total AMPK and α-SNAP at various time points following anisomycin incubation in cortical synaptoneurosomes. Data presented as mean +/- SEM. Values in all other groups were normalized to the 0 min group for the corresponding experiment and all the data points are represented as a fraction of the initial time point. n=2-3 animals per group.

**(C)** Representative immunoblot and its related quantification showing the reduction in the α-SNAP level on SNAP siRNA treatment compared to scrambled siRNA. Data presented as mean +/- SEM as a fraction of the scrambled siRNA group. Values in SNAP siRNA group were normalized to the control siRNA group for the corresponding experiment. *p<0.05, n=5 30mm dishes from 3 independent platings, One-sample t Test.

**(D)** Representative images depicting intracellular Ca^2+^ levels through Fluo8 fluorescence (Pseudo-colored) of a cortical neuron before stimulation, 15 sec (immediate) after NMDA (20μM) treatment, 300 sec (delayed) after NMDA treatment and after ionomycin treatment in the presence of 10mM Ca^2+^ and in the presence of NMDAR antagonist D-AP5 and NMDAR channel blocker MK-801. Scale bar 20μm.

**(E)** Representative immunoblots describing changes in phospho-eEF2 and total-eEF2 levels at various time points after NMDA (40μM) treatment to cortical synaptoneurosomes in the presence or absence of extracellular Ca^2+^. Note, in each case phospho and total eEF2 levels were normalized individually to their respective tubulin levels before calculating the phospho/total ratio.

**(F)** Line graph showing the average value for synaptic phospho/total eEF2 at various time points after NMDA treatment in the presence or absence of Ca^2+^. Data presented as mean +/- SEM. Values in all other groups were normalized to the 0 min group for the corresponding experiment and all the data points are represented as a fraction of the initial time point. *p<0.05, **p<0.01, n≥ 4 animals per group. Two Way ANOVA followed by Bonferroni’s multiple comparison test.

**(G)** Bar Graph representing phospho/total eEF2 ratio on 5 minutes of NMDA stimulation in cortical synaptoneurosomes in the presence or absence of extracellular Mg^2+^. Data presented as mean ± SEM as a fraction of their respective basal condition. n=4 animals per group.

**(H)** Representative images showing phospho-eEF2 immunolabeling (Pseudo-colored), newly synthesized protein through FUNCAT metabolic labeling and MAP2B immunolabeling in cortical neurons on AICAR (1mM) treatment and Compound C (10μM) treatment for 1 hour. MAP2B immunolabeling was used for identifying neurons and intensity was used for normalization. Scale bar 10μm.

Conflict of Interest……………………………………………………………………………………………………….34

References…………………………………………………………………………………………………………………….34

Figure Legends……………………………………………………………………………………………………………….46
