## Supplementary figures and images for "Distinct Regulation of Bioenergetics and Translation by Group I mGluR and NMDAR"

### Supplementary Figure 1

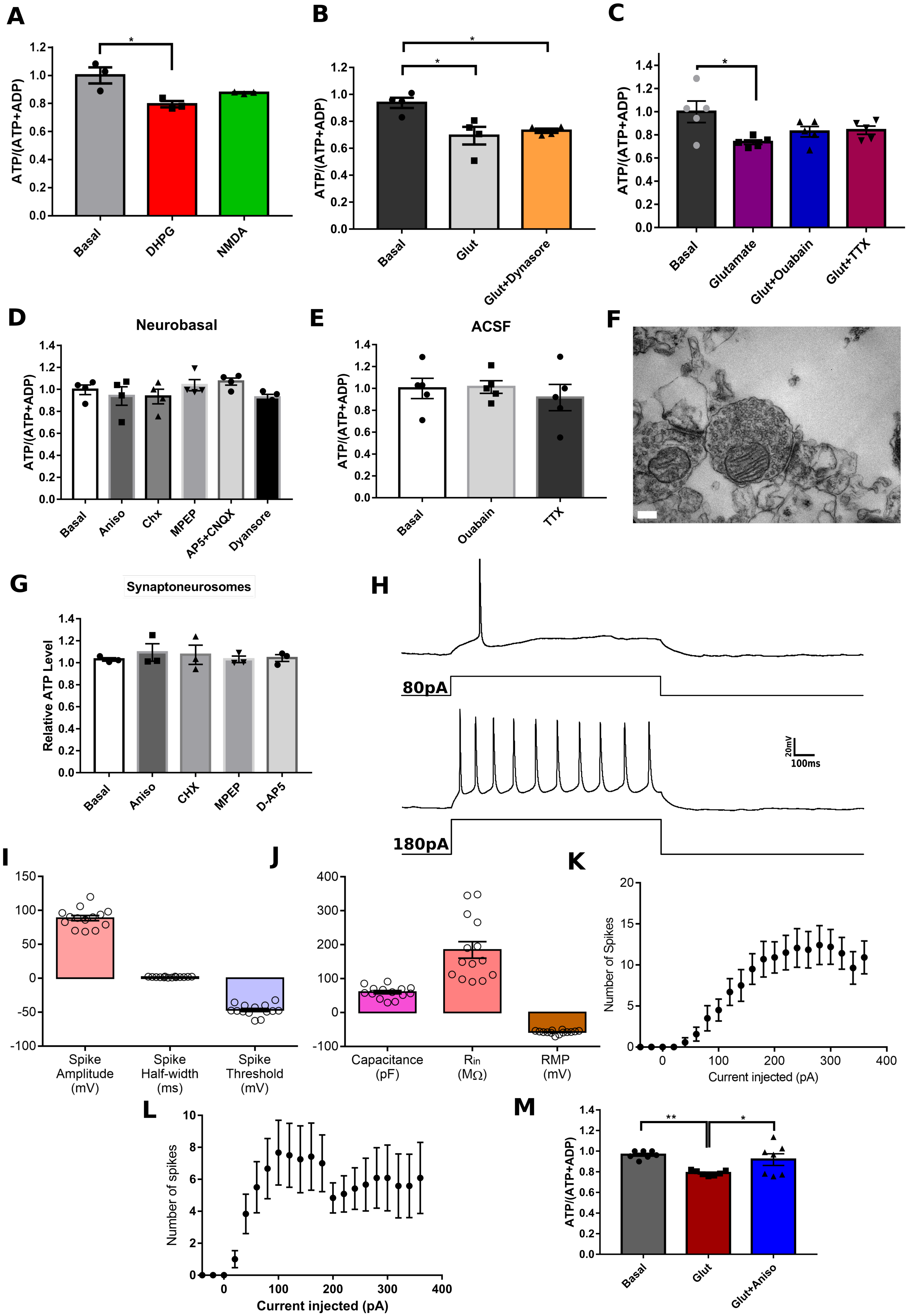

### Supplementary Figure 2

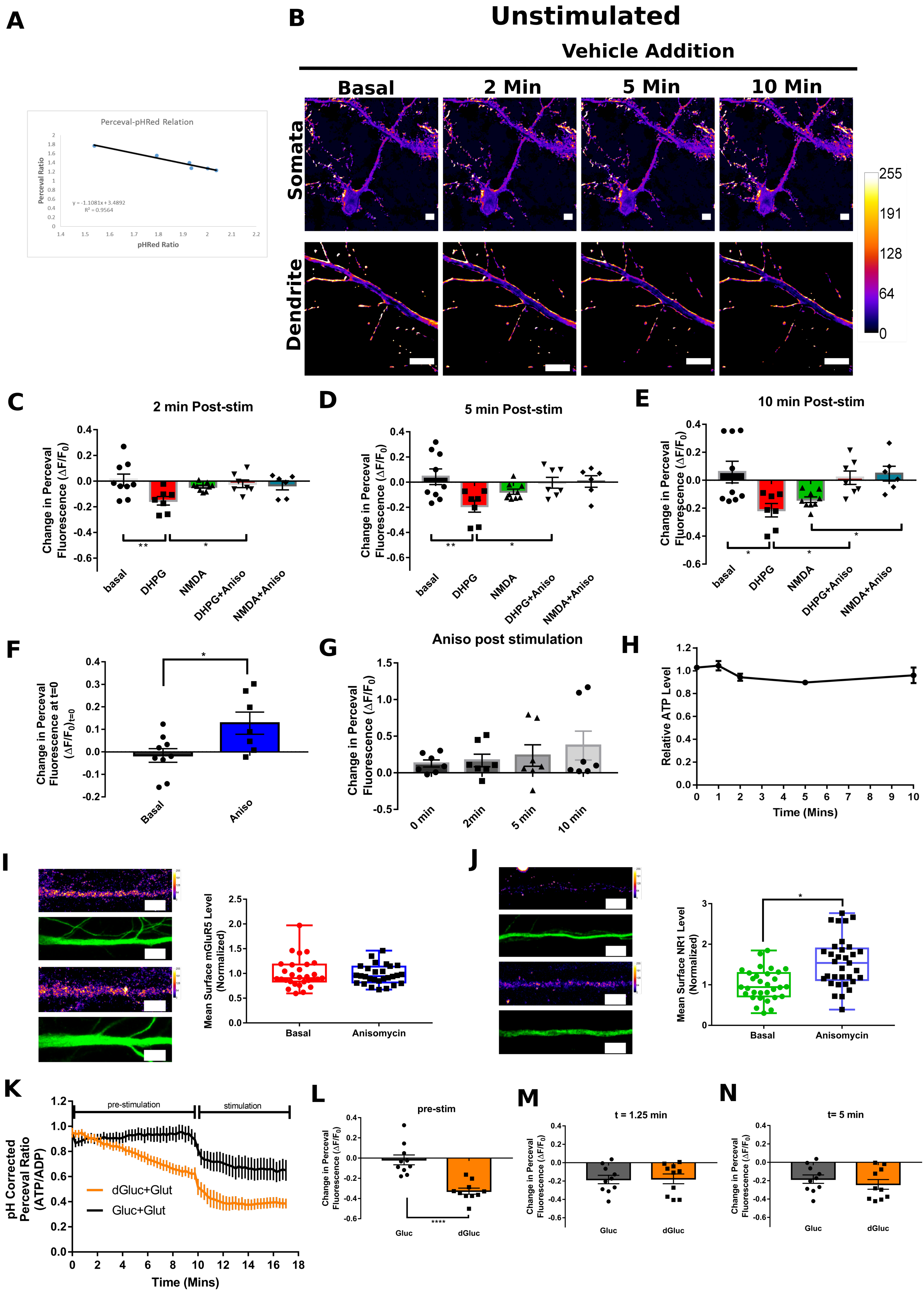

### Supplementary Figure 3

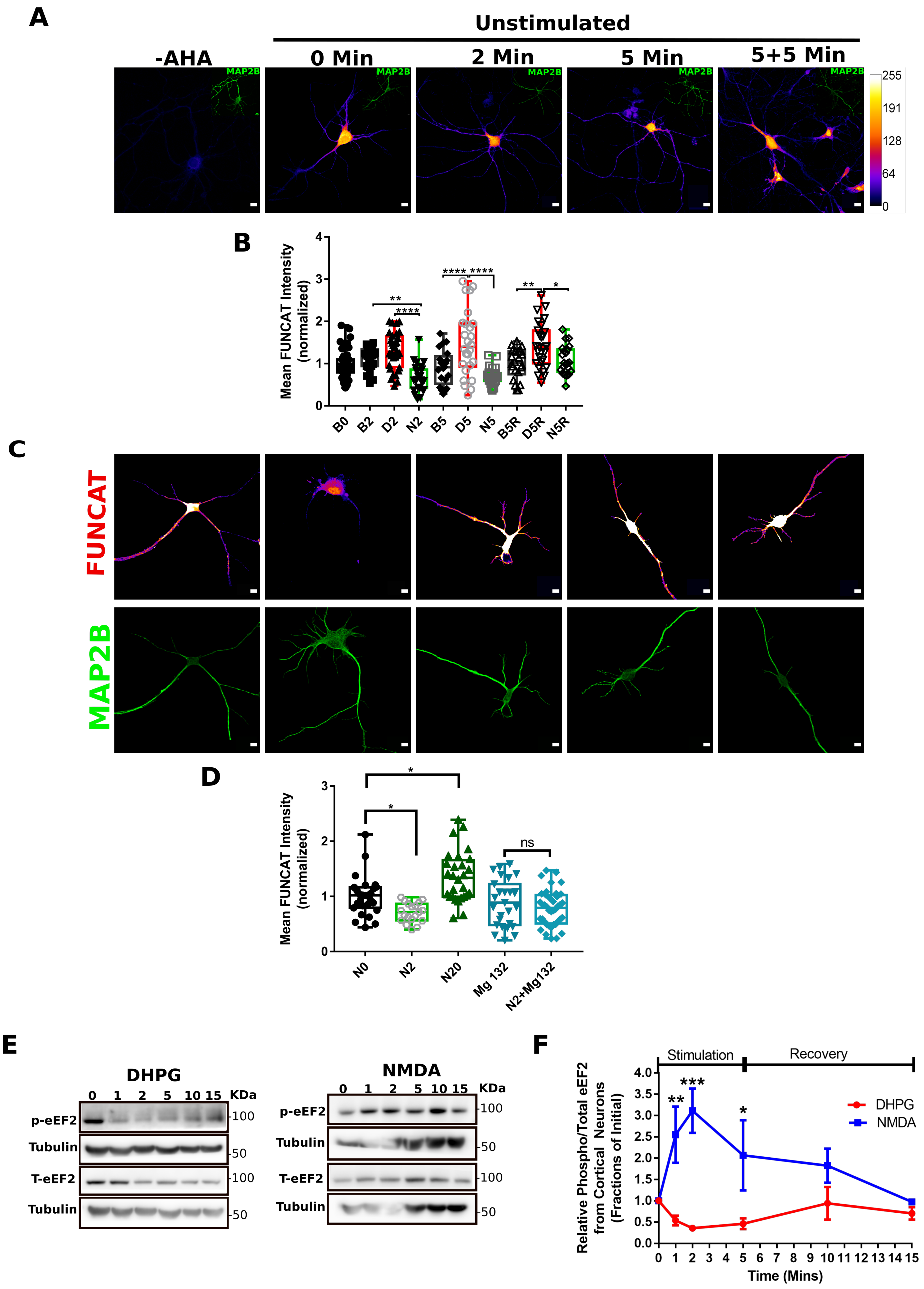

### Supplementary Figure 4

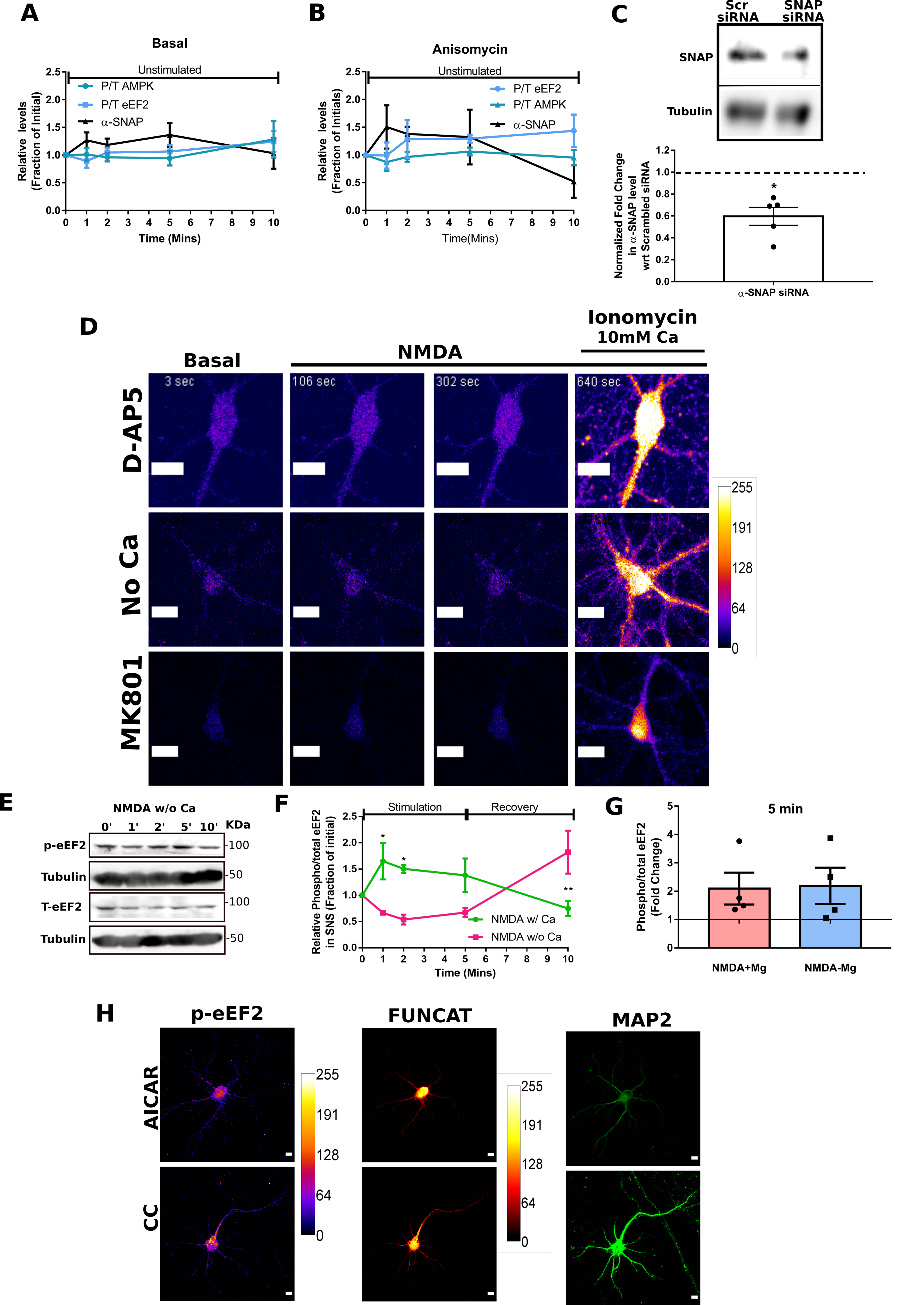
